# Behaviour of pyrethroid resistant Anopheles gambiae at the interface of two dual active-ingredient bed nets, assessed by room-scale infrared video tracking

**DOI:** 10.1101/2022.07.20.500766

**Authors:** K. Gleave, A. Guy, F. Mechan, M. Emery, A. Murphy, V. Voloshin, C. E. Towers, D. Towers, H. Ranson, G. M. Foster, P. J. McCall

## Abstract

**Background:** The success of Insecticide Treated Bednets (ITNs) for malaria vector control in Africa relies on the behaviour of the major malaria vectors, Anopheles species. Research into mosquito behavioural traits influencing the performance of ITNs has focused largely on time or location of biting. Here we investigated less tractable behaviours including timings of net contact, willingness to refeed and longevity post exposure to two next-generation nets, PermaNet® 3.0 (P3) and Interceptor® G2 (IG2) in comparison with a standard pyrethroid only net (Olyset (OL)) and an untreated net.

**Methods:** Susceptible and resistant Anopheles gambiae mosquitoes were exposed to the nets with a human volunteer host in a room scale assay. Mosquito movements were tracked for two hours using an infrared video system, collecting flight trajectory, spatial position and net contact data. Post-assay, mosquitoes were monitored for a range of sublethal insecticide effects.

**Results:** OL, P3 and IG2 all killed over 90% of susceptible mosquitoes 24 hours after exposure, but this effect was not seen with resistant mosquitoes where mortality ranged from 16% to 72%. Total mosquito activity was higher around untreated nets than ITNs. There was no difference in total activity, the number, or duration, of net contact, between any mosquito strain, with similar behaviours recorded in susceptible and resistant strains at all ITNs. Net contact was focussed predominantly on the roof for all bednets. We observed a steep decay in activity for both susceptible strains when P3 and OL were present and with IG2 for one of the two susceptible strains. All treated nets reduced the willingness of resistant strains to re-feed when offered blood one-hour post-exposure, with a more pronounced effect seen with P3 and OL than IG2.

**Conclusion:** Results indicate that the effects of ITNs on mosquito behaviour are consistent, with no major differences in responses between strains of different pyrethroid susceptibility.

## Background

Resistance to insecticides has emerged in mosquitoes across the globe and threatens the future use of insecticides to control many vector-borne diseases. The most effective malaria control method in Africa, where the vast majority of malaria cases occur, is the widespread use of insecticide-treated nets (ITNs) (Pryce et al., 2018). The first generation of ITNs use fast-acting pyrethroids, and pyrethroid resistance has spread at an alarming rate through Anopheles populations in Africa (Hancock et al., 2020; Hemingway, 2017; Ranson & Lissenden, 2016) reducing ITN efficacy (Churcher et al., 2016). Several types of ‘next-generation ITNs’ are now available and used in many malaria-endemic countries; these all contain pyrethroids plus an additional active ingredient (AI) with a different mode of action (MoA). Currently, the most widely used next-generation nets are pyrethroid-piperonyl butoxide nets (pyrethroid-PBO nets); PBO increases the potency of pyrethroids by blocking enzymes that break down insecticides. In 2021, pyrethroid-PBO nets constituted 42.8% of the nets distributed in Sub-Saharan Africa with public funds (The Alliance for Malaria Prevention, 2022). Recent clinical trials of ITNs with two insecticides (Interceptor G2®, BASF, containing a pyrethroid plus the pyrrole insecticide chlorfenapyr) (Mosha et al., 2022) or containing pyrethroid plus pyriproxyfen (a chemical that sterilises female adults) (Tiono et al., 2018) have shown improved clinical outcomes over standard ITNs. However, improved epidemiological outcomes have only been demonstrated in a single setting with pyriproxyfen nets showing no improved public health value over standard ITNs in the Tanzanian trial (Mosha et al, 2022). Further evidence of their efficacy in different ecological and epidemiological environments is needed prior to national or global policy changes.

The success of ITNs relies predominantly on the daily behaviour of the major malaria vectors in Africa, where Anopheles species are largely anthropophagic, endophagic, endophilic and feed during the night when people are more likely to be underneath their bed nets (Pates & Curtis, 2005; Killeen et al., 2006). Multiple types of mosquito behavioural alterations in response to widespread ITN use at the population level could decrease their efficacy (Gatton et al., 2013; Killeen, 2014), and several examples of this behavioural resistance have been described after multiple years of net use. For example following a mass ITN distribution programme in Benin, An. funestus have shown a shift in biting time from a peak late at night to early morning when people emerge from their protective ITNs (Moiroux et al., 2012). Monitoring these population changes induced by widespread deployment of ITNs, or any other vector control tool, is essential to explain and predict their epidemiological impact. Indeed, modelling studies have indicated that behavioural resistance and physiological resistance (caused, for example, by target site modifications or enhanced detoxification) could be equally detrimental to the efficacy of ITNs (Gatton et al., 2013). Therefore, surveillance of vector behaviour is an essential component of resistance management programmes.

In addition to population surveillance, critical insights into the behaviour of mosquitoes in response to ITNs can be gained by laboratory and semi-field studies that quantify important traits This includes net contact time and blood-feeding volumes and relates these to key endpoints such as longevity and reproductive outputs. Performing these tests on mosquito populations with different levels, and mechanisms of pyrethroid resistance, may inform predictions on the efficacy of standard and next-generation ITNs in different environments. Standard WHO assays, designed to measure the performance of a single, fast-acting insecticide in ITNs (i.e pyrethroids) are not suitable for measuring the impact of combining AIs with differing MoAs and endpoints. We have therefore been developing and evaluating a series of benchtop and room-scale assays to record mosquito responses to a more diverse range of ITNs.

The ‘baited box’ assay allows for close-range observation of mosquitoes attempting to take a blood meal through an ITN, with results from Hughes et al., reporting that the accumulated duration of net contact by Anopheles gambiae was 50% lower on ITNs compared to untreated nets, with no difference in contact duration between susceptible and resistant mosquitoes (Hughes et al., (2020). Benchtop tests are undoubtedly informative, but the impacts of ITNs extend beyond the close range captured in these assays. Parker et al., (2015, 2017) used an infrared tracking system to characterise mosquito behaviour at mid-range, i.e. host-seeking events around an entire human-baited PermaNet® 2.0 bed net (Vestergaard Sarl), from room entry to arrival at the ITN. The initial behaviour of insecticide-susceptible An. gambiae and wild An. arabiensis did not differ between an untreated or pyrethroid ITN; mosquitoes continued to respond to the host without any evidence of repellency until they contacted the insecticide on the net surface. After this time, activity decayed rapidly, reaching zero after around 30 minutes, demonstrating the highly efficient rapid action of pyrethroid-treated ITNs. Here we apply this method to studying the behaviour of insecticide-resistant mosquitoes to next-generation bed nets to gain initial insights into the utility of this method in comparing responses between mosquito populations and net types.

This study investigated the mosquito response to two next-generation nets, PermaNet® 3.0 (Vestergaard Sarl) and Interceptor® G2 (BASF AGRO B.V Arnhem [NL] Freienbach Branch) performed in comparison with a standard pyrethroid only ITN (Olyset Net, Sumitomo Chemical Co., Ltd) and an untreated net, as measured by impacts on both pyrethroid susceptible and resistant mosquitoes. This study also sought evidence for any altered behaviours during host-seeking at the net which may be attributed to the new nets.

## Materials and Methods

Mosquitoes from two insecticide-susceptible (Kisumu and N’gousso) and two insecticide-resistant (VK7 and Banfora) An. gambiae s.l strains were maintained under standard insectary-controlled conditions (27°C ± 2°C, and 80% relative humidity (RH)) at the Liverpool School of Tropical Medicine (LSTM). Susceptible An. gambiae s.s Kisumu colony originates from Kenya (Shute, 1956) and has been maintained in colony since 1975. An. coluzzii N’gousso was colonised from Cameroon in 2006 (Harris et al., 2010). An. coluzzii VK7 and Banfora strains originated from Burkina Faso, have been reared at LSTM since 2014 and 2015, respectively, and are highly resistant to pyrethroids with susceptibility only partially restored by PBO pre-exposure (Williams et al., 2019, 2022). The VK7 strain is fixed for the knockdown resistant (Kdr) 995F allele in the voltage-gated sodium channel (Vgsc), whereas the Banfora strain has a more complex set of Vgsc mutants (Ingham et al., 2021). Both strains have elevated cytochrome P450 expression, but additional resistance mechanisms are present in the Banfora strain including an increased respiratory rate (Ingham et al., 2021). All mosquitoes were reared under an altered 12:12 light/dark cycle to allow for testing to be conducted during the ‘night’ phase of the circadian rhythm.

The ITNs used are shown in Table 1. Nets were obtained directly from the manufacturer, aired at room temperature for four weeks prior to testing and then adjusted in size to fit the custom-made bed net frame, ensuring maximum visualisation of mosquito activity. A single net was used for each treatment, each stored at 4°C between testing replicates and acclimatised at 27±2°C and 70±10% humidity for at least one hour prior to testing.

**Table 1.** Insecticide treated nets used in room scale tracking assays.

| *Net type* | *Specification* | *Manufacturer* |
| --- | --- | --- |
| Polyester control | Untreated | Bayer AG, Leverkusen, Germany |
| Olyset Net (OL) | 150 denier polyethylene net incorporated with permethrin at 800 mg/m<sup>2</sup> | Sumitomo Chemical Company, Tokyo, Japan |
| PermaNet 3.0 (P3) | Roof: 100 denier polyethylene net incorporated with deltamethrin at 120mg/m<sup>2</sup> and PBO at 750mg/m<sup>2</sup>, Sides: 75 denier polyethylene net with deltamethrin at 84mg/m<sup>2</sup> | Vestergaard Sarl, South Africa |
| Interceptor G2 (IG2) | 75 denier polyester net coated with alphacypermethrin at 100mg/m<sup>2</sup> and chlorfenapyr at 200 mg/m<sup>2</sup> | BASF AGRO B.V Arnhem (NL), Germany |

All experiments required a human volunteer to act as bait under the net. Volunteers were asked to wear light clothing, not to wear any strong scented products and not to bathe for at least four hours prior to testing. During the experiment, volunteers were asked to lie as motionless as possible, while still being comfortable. To control for any effect of body positioning, volunteer orientation was randomly assigned either with head or feet nearest to the mosquito release point.

A total of 25, three-to-five-day old un-fed female mosquitoes were used per test replicate, as per Parker 2015 (Parker et al 2015). Mosquito access to 10% sugar solution was removed by 16:00 the day prior to testing and replaced with distilled water; this was removed three hours prior to testing.

### Experimental set-up

All experiments were performed in the LSTM Accelerator building, using a custom built free-flight testing room (7m x 4.8m in area, 2.5m high) which is climate controlled (27±2°C and 70% ±10% RH), while recording is operated from an adjacent room. Assays were performed during the afternoon to coincide with the ‘night’ phase of the mosquito’s circadian rhythm when they would be host-seeking in the wild. Frames made of carbon rods with roofs tilted towards the recording equipment were constructed for each bed net type to allow accurate observations of mosquito activity (dimensions: front height 45cm, rear height 75cm, roof width 90cm, roof length 180cm).

Mosquitoes were placed into a holding cup one hour prior to testing to acclimatise within the testing room. The cup was attached to a long cord allowing mosquitoes to be released remotely by the operator outside the tracking room. Fifteen minutes before the test began the volunteer entered the ITN; to start the test, the release cord was pulled. After two-hour recording, free flying and knocked down mosquitoes were collected using a HEPA filter mouth aspirator (John. W. Hock, USA) to avoid any insect damage and placed into a fresh collection cup. Mortality was recorded at 24 hours, with all mosquitoes individually monitored for sub-lethal insecticide effects (see below).

ITN treatments were changed approximately every three weeks and the testing room decontaminated between each ITN type, using 5% Decon90 solution (Decon Laboratories Conway Street, UK), followed by two water washes and a final wash with 70% ethanol. World Health Organisation (WHO) cone tests (World Health Organization, 2006) using susceptible An. gambiae were performed on the walls 24hours after decontamination for quality control (QC). During testing, no WHO cone assays resulted in >20% mortality, therefore all cleaning procedures were considered to pass the QC process.

### Mosquito Tracking

Mosquitoes were tracked using paired identical recording systems, positioned 1050 mm apart and consisting of the following: each recording system used one camera (12 MPixel Ximea CB120RG-CM with a 14mm focal length lens), aligned with a single Fresnel lens (1400 x 1050mm and 3mm thick, 1.2m focal length; NTKJ Co., Ltd, Japan) placed approximately 12100 mm away. Cameras recorded with an exposure time of 5ms and −3.5 dB gain with a lens aperture of F#8.0 (Voloshin et al., 2020). As experiments were carried out in the dark, infrared light was provided using custom ring light sources constructed by colleagues at Warwick university (12 OSRAM™ SFH 4235 infrared LEDs with a peak wavelength of 850nm) which illuminated the total recording volume of 2 x 2 x 1.4m. To reflect light back towards the cameras a custom designed Retroreflective screen (2.4 x 2.1 m, material: 3M™ Scotchlite™ High Gain Reflective Sheeting 7610) was placed 2m from the Fresnel lenses, with the bed and ITN placed in between both. The reflected light is focused by the Fresnel lens and forms a telecentric lens pair with an imaging optic mounted on the camera which allows illumination and imaging to occur from one side of the experimental set up. More information on signal processing can be found in Voloshin et al., (2020). Recordings were captured for both cameras over the two-hour assay using StreamPix recording software (StreamPix V7, Norpix, Montreal, Canada) at 50 frames per second (fps) onto a Windows PC (Intel® Xeon® Silver 4114 CPU 2.20 GHz, 24 Gigabytes RAM, Windows 10 Pro; 12 configured into 2 RAID arrays of 24 Terabytes each, at 1 array per camera.

### Video analysis

All video analysis was carried out using bespoke software written in Matlab (Mathworks) developed by collaborators at Warwick University (Angarita-Jaimes et al., 2016). Video segmentation, then compression to .mp4 files was performed before all videos were manually reviewed and cleaned to remove false tracks and human movement using ‘Seq File Processing’ software. Data extracted includes trajectory duration, distance travelled, the number, duration and location of contacts with the bed net, time to first contact and track velocity, all of which have been previously described by Parker et al., (2015). Additional track joining and the deletion of false tracks created by volunteer and camera noise was performed in ‘Post Processing’ along with categorising activity into behavioural modes using existing quantification algorithms (Table 2) and dividing the field of view into 10 distinct regions to quantify net contact location and duration at 10 different regions of the bed net. Since multiple mosquitoes were released into the room in all tests, tracking individual mosquitoes was not possible, hence analysis was performed on flight tracks with each track from entry into and exit out of the field of view analysed separately. One flight track could consist of three different behavioural modes (visiting, bouncing and resting as they all involve net contact), upon which the time spent in each mode were recorded separately.

**Table 2.**
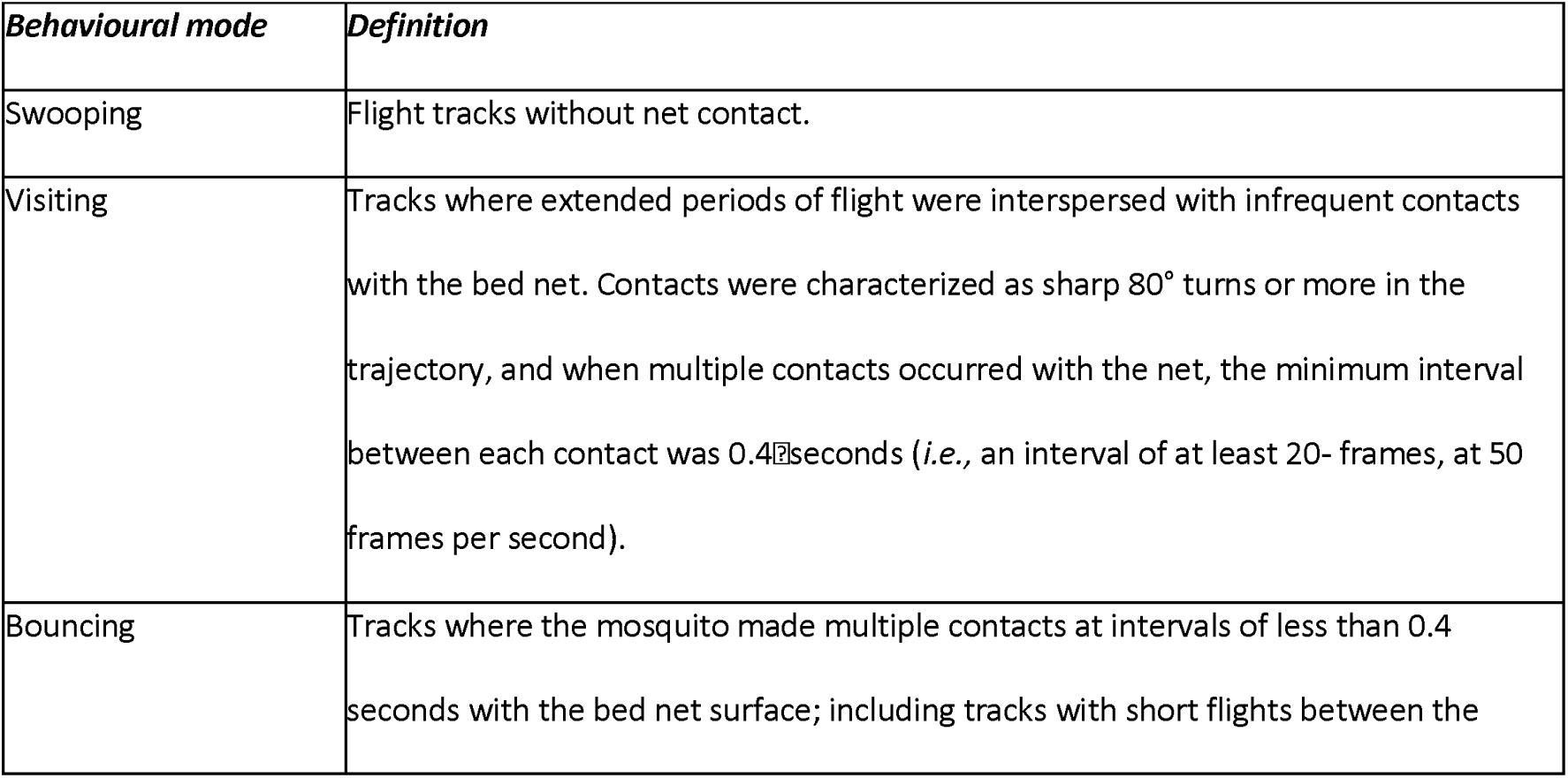
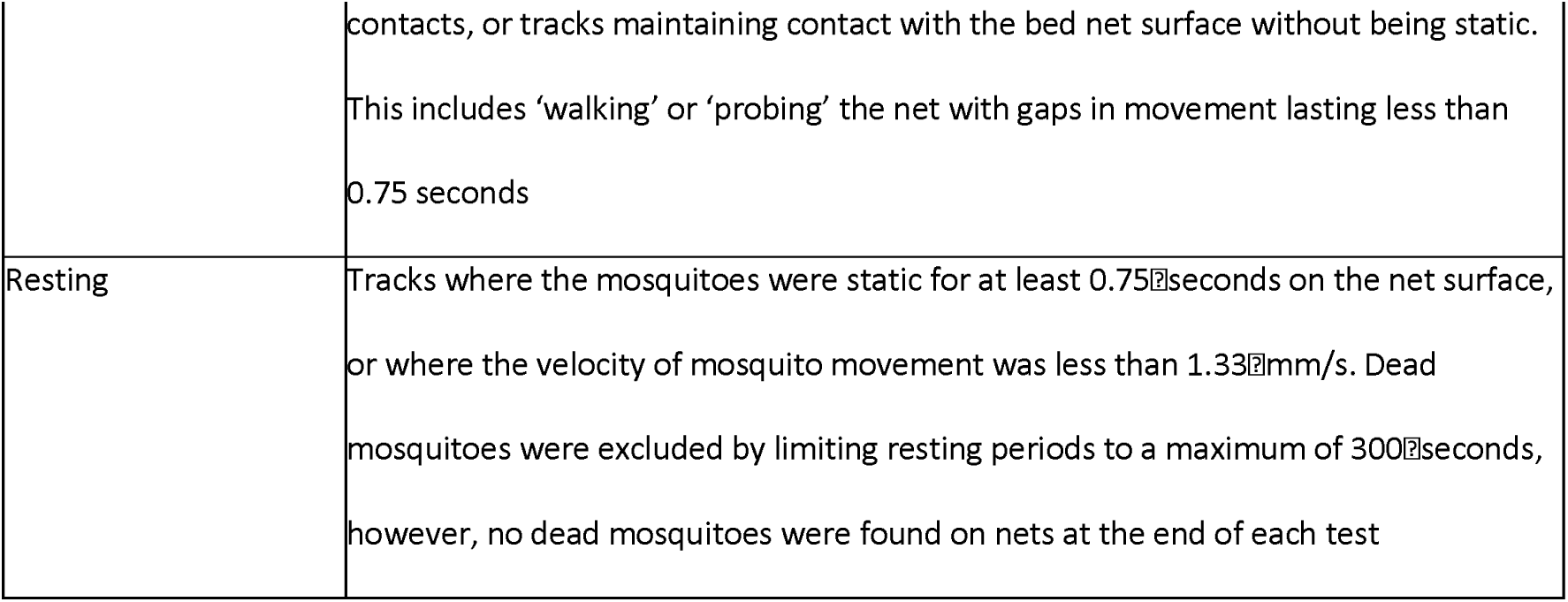
Definition of mosquito behavioural modes (adapted from (Parker et al., 2017).

### Sub-lethal pipeline

The methods for sub-lethal pipeline monitoring have been previously described in Hughes et al., (in press). After each tracking assay, the following were measured for each individual mosquito: 24hour mortality, willingness to feed at 60 minutes, or 24hours (by exposure to the arm of a human volunteer), longevity and wing length (Figure 1).

**Figure 1.**
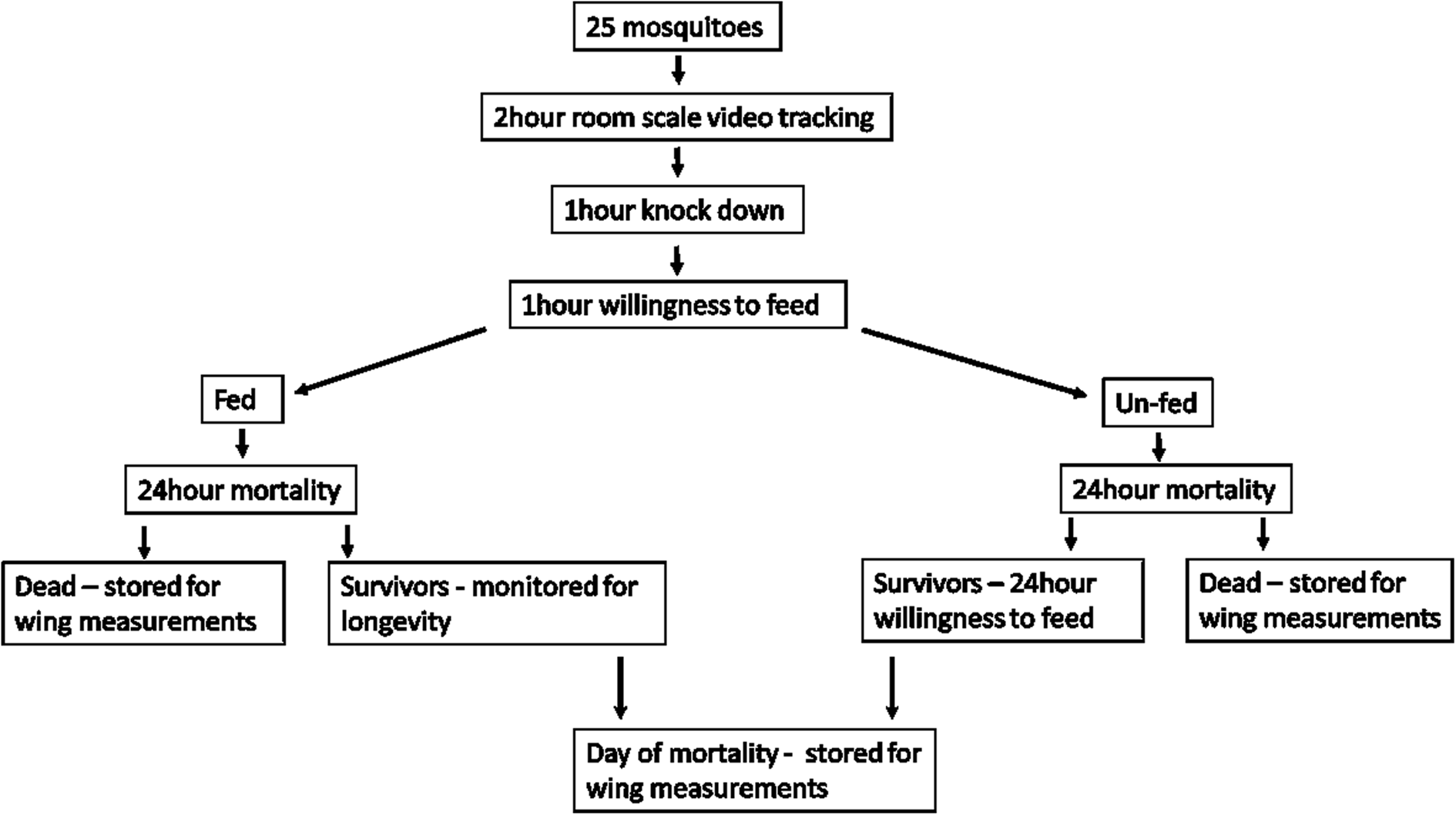
Measured sub-lethal pipeline outcomes per room scale video tracking assay.

### Data analysis

A sample size for comparing net contact times at three different ITNs was calculated using the mean difference in net contact time for a single strain between untreated and treated nets generated in an earlier study in the statistical program R (Version, 1.1.463, R Core Team 2019), and using the phia (Rosario-Martinez, 2015) and pwr (Champely, 2017) packages. With a significance level of 0.05 that gives at least a power of 90%, a minimum sample size was determined inflating the sample size with 30% to adjust for any potential confounding factors. A common standard deviation was assumed for all groups used was 562.14s and mean differences were 4.54 time reduction in PermaNet 2.0 arm compared to untreated, and 5 times reduction in Olyset group compared to untreated (obtained from the previous study based on the ANOVA or t-test (Parker et al., 2017)). A total of 6 replicates per strain and treatment was the minimum requirement determined to compare net contact times at different ITNs. This sample size does not account for the correlation of the measurements from the same volunteer, although this correlation may still exist.

### ITN bioefficacy and mosquito longevity

Bioefficacy of nets was assessed through measuring mosquito mortality post-exposure. Mosquitoes were transferred to individual falcon tubes, provided with a source of 10% sugar water and mortality measured daily until all mosquitoes had died.

### Quantifying mosquito activity and behaviour

Total activity per strain (seconds of movement), per net treatment was calculated as the sum of all mosquito activity, regardless of behavioural mode and binned into 5-minute intervals for analysis. Further analyses were performed using the total activity stratified into the four described behavioural modes (swooping, visiting, resting and bouncing).

### Defining and quantifying mosquito contact with the bed net interface

Total contact number and total contact duration with a net was calculated from the sum of all contacts obtained from visits, bounces or resting tracks. Total duration of contact in the first 10 minutes of the assay was calculated as a percentage from overall contact duration along with an average of mosquito duration. As it was not possible to determine individual mosquito contact, we calculated the possible minimum and maximum values of net contact as in Parker et al., (2015): for the maximum value, total contact duration was divided by the maximum number of mosquitoes seen simultaneously contacting the net in any one frame of the recording; the minimum value assumed that all 25 mosquitoes released into the assay responded at the same time.

### Determination of contact location

The recording field of view was divided into 16 regions using previously described software (Angarita-Jaimes et al., 2016). Ten of these regions were on the net surface; six on top of the bed net, two on the front of the net and one at either side.

### Speed around the bed nets

Flight speed was analysed using whole swooping tracks around the bed nets to investigate any changes in mosquito flight.

### Mosquito activity decay over the 2hour assay

Exponential decay modelling was considered for analysis of activity over time, as reported previously by Parker et al., (2015) but many of the test replicates violated the equation constraints, so an alternative method was used whereby total activity in the first 5 minutes of recording was subtracted from total activity in the final 5 minutes of recording. A negative value indicated that activity decayed over time and a positive value represented an increase in activity between the two timepoints.

### Determining willingness to refeed and mosquito size

Wing length was used as an estimate for mosquito body size and to control for potential size differences between cohorts. The right wing was removed, and an image taken using GXCAM ECLIPSE Wi-Fi camera attached to a GX Stereo microscope (GT Vision Ltd). The length of the wing was measured from the axial vein to the distal end of the R1 vein using GXCAM software (GXCAM Ver.6.7).

To assess any effects of sub-lethal insecticide exposure, mosquitoes were offered a blood meal at 1-hour post-exposure and longevity measured. Blood feeding inhibition was calculated by considering all mosquitoes in each replicate and assessing whether they were able to take a blood meal or not.

### Statistical analysis

Statistical analysis was performed used Prism 6 (GraphPad) and R (Version, 1.1.463, R Core Team 2019). 24hour mortality was assessed using t-tests for the comparison of observed means, and mosquito longevity was analysed using Kaplan Meir Long-rank (Mantel-Cox) tests. Shapiro-Wilk tests were carried out on all activity data to check for normality. Total activity was analysed used Welch’s ANOVA as we did not assume that all groups sampled were from populations with equal variance. Generalised linear models (GLMs) with normal probability distribution were used to analyse pairwise comparisons of mosquito strain and net type for: behavioural mode, contact number, contact duration, duration of contact in first 10 minutes, average contact duration, swooping speed, activity decay, willingness to refeed and wing length. Post-hoc analysis used the Tukey method of adjustment for comparing a family of four estimates. We used a binomial GLM to look for any interactions that might explain a relationship between net contact duration and mortality, however the model showed that there was no interaction between net type and contact duration or strain and contact duration. We used a GLM to investigate the relationship between mosquito wing size and blood feeding success, considering interactions with mosquito strain and net type. For all statistical comparisons, the α threshold used was 0.05. Unless stated otherwise, 95% confidence intervals are reported.

### Ethical permission

With no infection risk and no exposure to untested chemicals, the procedures involved in generating these data results did not require clearance by LSTM Research Ethics Committee. We obtained written consent from all volunteers.

## Results

A total of 1690 mosquitoes were tested across 73 assays, with 18 different volunteers being used as a human ‘bait’. The total number of replicates performed for each strain and treatment are shown in Table 3. It was not possible to reach the target replicate number of six for all strain and net treatment combinations because several video files were corrupted during a computer failure resulting in missing videos, time constraints due to national COVID-19 restrictions and the LSTM Banfora colony which lost its high level of resistance before PermaNet 3.0 and Interceptor G2 replicates could be completed. All room scale recordings were completed between June 2019 and February 2020.

**Table 3.** Total number of replicates performed per ITN, per mosquito strain. (UT = untreated net, OL = Olyset Net, P3 = PermaNet 3.0, IG2 = Interceptor G2)

| *Strain* | *Recording dates* | *Long-Lasting Insecticidal Nets* |  |  |  |
| --- | --- | --- | --- | --- | --- |
|  |  | *UT* | *OL* | *P3* | *IG2* |
| **Kisumu** | **Jun 2019 –Jan 2020** | 5 | 6 | 6 | 6 |
| **N'gousso** | **Jun 2019 – Nov 2019** | 4 | 6 | 2 | 6 |
| **VK7** | **Jun 2019 – Feb 2020** | 4 | 5 | 5 | 5 |
| **Banfora** | **Jul 2019 – Dec 2020** | 4 | 6 | 3 | 0 |

### Mosquito survival

#### Bioefficacy

Mortality at 24h after the two-hour room scale tracking assay on untreated net (UT) was below 20% for all strains (Figure 2). OL, P3 and IG2 all killed more than 90% of susceptible strains within 24hours. Mortality rates at 24hours were significantly lower for resistant VK7 and Banfora strains with OL, P3 and IG2 nets (Figure 2) (Additional Table 1) compared to susceptible strains Kisumu and N’gousso (OL: VK7 v Kisumu p<0.0001, VK7 v N’gousso p<0.0001, Banfora v Kisumu p=0.0013, Banfora v N’gousso p=0.0014; P3: VK7 v Kisumu p=0.0042, N’gousso v VK7 p=0.0903, N’gousso v Banfora p=0.0602 Banfora v Kisumu p=0.0007; IG2: VK7 v Kisumu p<0.0001, VK7 v N’gousso p<0.0001) (Additional Table 2). Note that the N’gousso results derive from only 2 test repeats, which may account for the non-significant P-values, despite the differences in mean mortalities. The highest 24hour mortality observed for VK7 strain was following P3 tests, which was significantly higher than that of OL (p=0.0009) and IG2 (p<0.0001). There was no significant difference in mortality rates between OL and IG2. Twenty-four-hour mortalities of the Banfora strain ranged between 45.34% and 72.38% and were not significantly different between ITNs.

**Figure 2.**
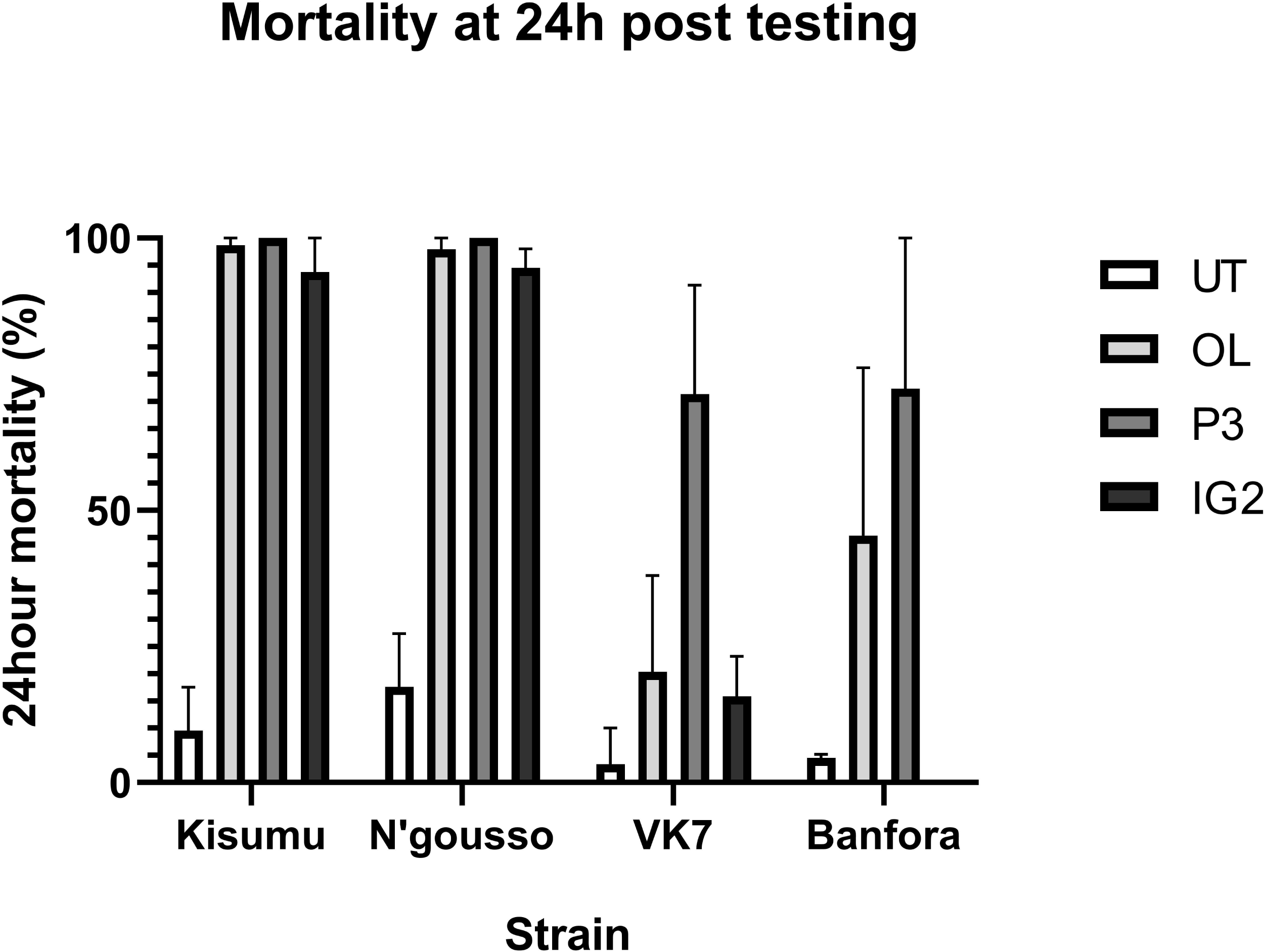
Mean mortality of two susceptible (Kisumu, and N’gousso) and two resistant (VK7 and Banfora) *Anopheles gambiae strains* at 24h after a two-hour exposure during room scale tracking to untreated net (UT), Olyset Net (OL), PermaNet 3.0 (P3) and Interceptor G2 (IG2) with 95% Confidence Intervals.

Cumulative mortality rates 72hr after exposure to IG2 (containing the slower acting pyrrole insecticide chlorfenapyr) were lower in VK7 than in both susceptible strains (VK7 25.25%, 95% CI 10.29, 40.21]; Kisumu 95.91%, 95% CI [86.91, 100]; N’gousso 98.86%, 95% CI [95.25, 100]; VK7 v Kisumu t(8)= 9.28, p<0.0001; VK7 v N’gousso t(8)= 10.04, p<0.0001). Cumulative 72hr mortality for VK7 and Banfora after exposure to OL increased to 35.04% and 61.42% respectively, and after P3 exposure to 79.29% and 73.53% respectively. The increase in mortality between 24 and 72 hours seen after all ITN exposure was not significantly different to the increase seen in this time frame after exposure to UT nets for either resistant strains.

#### Longevity

For VK7, median survival time after IG2 exposure was identical to that recorded after UT exposure IG2 10days [95% CI 7.53, 12.48]; UT 10days [95% CI 8.23, 11.77]] with no significant difference in overall longevity [VK7 UT v IG2 p=0.2150]. For the same strain, median survival times following OL exposure was five days [95% CI 3.20, 6.80] and following P3 was one day [95% CI 0, 1]. In both resistant strains, P3 exposure had the largest impact in reducing longevity (VK7: UT v OL p=0.0198, UT v P3 p<0.0001; Banfora: UT v OL p=0.0026, UT v P3 p=0.0099) (Figure 3). Both resistant strains survived significantly longer after exposure to all three ITNs compared to the susceptible strains (Additional Table 3). The median survival time after exposure to UT nets varied between strains (Kisumu 7 days [95% CI 5.58, 8.33]; N’gousso 12 days [95% CI 10.25, 13.76]; VK7 10 days [95% CI 8.23, 11.77]; Banfora 8 days [95% CI 6.49, 9,51]).

**Figure 3.**
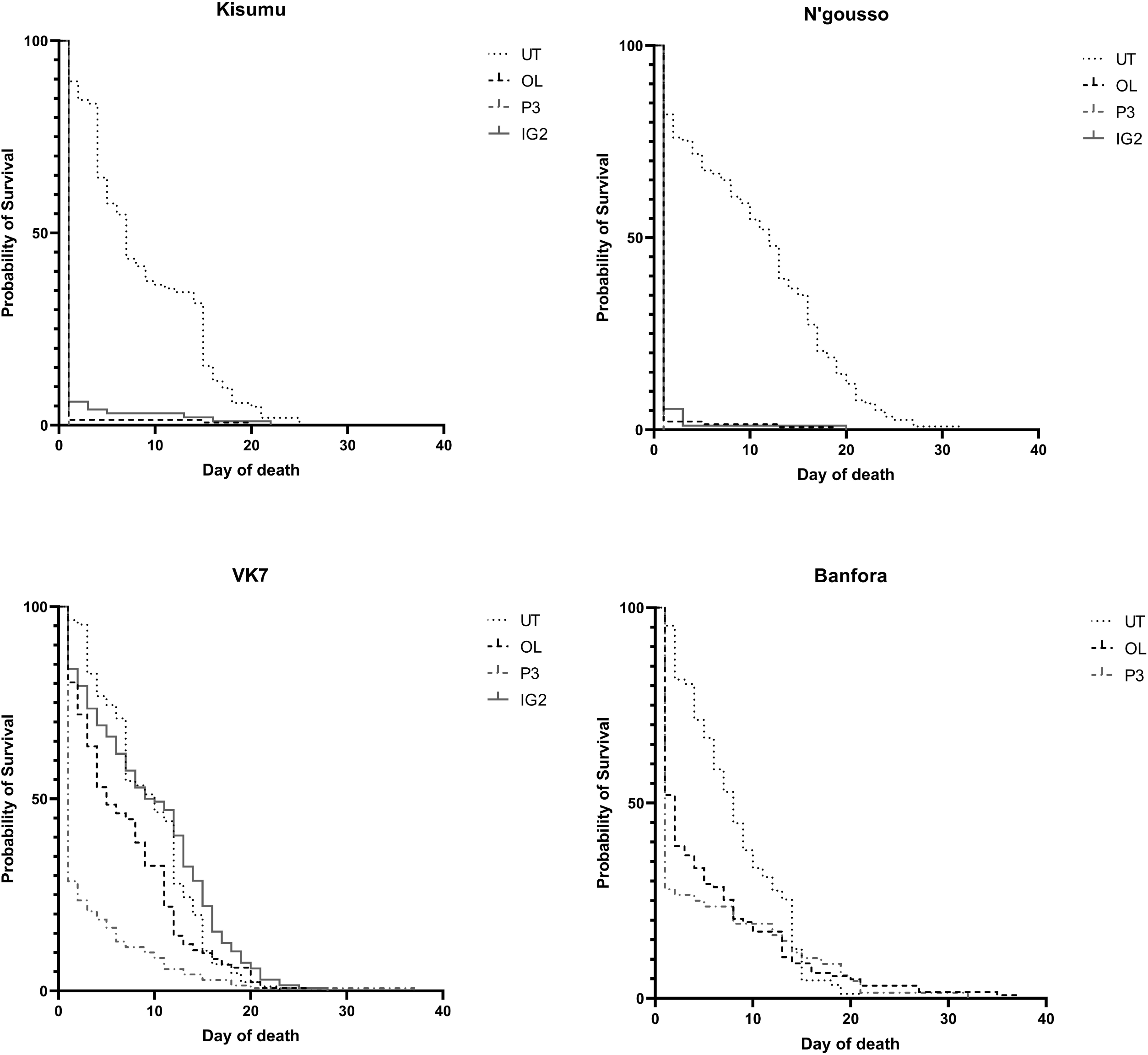
Survival curves for susceptible (Kisumu and N’gousso) and resistant (VK7 and Banfora) Anopheles gambiae after exposure in the room scale tracking room to either untreated net (UT), Olyset Net (OL), PermaNet 3.0 (P3) or Interceptor G2 (IG2). Day 0 is day of exposure.

### Mosquito activity and behaviour

#### Total activity and behavioural mode

Figure 4 shows mean total mosquito activity for each strain and net combination, across a two-hour recording, with activity separated into the four distinct behavioural modes: swooping, visiting, bouncing or resting defined by Parker et al (2015). Across all treatments, flight track length ranged from 2.5mm to 20,249mm and track duration ranged from 0.08 seconds to 1,010 seconds. For all four strains, total activity was significantly longer at an UT net than at any of the three ITNs (Kisumu Welch’s F(3.0, 8.71)=44.44, p<0.0001; N’gousso Welch’s F(3.0, 3.59)=24.15, p=0.0074; VK7 Welch’s F(3.0, 7.27)=20.82, p=0.0006; Banfora Welch’s F(2.0, 5.29)=32.17, p=0.0011). Comparing net types showed no significant differences in total activity between any of the strains (UT Welch’s F(3.0, 6.90)=3.94, p=0.0626; OL Welch’s F(3.0, 9.38)=2.21, p=0.1543; P3 Welch’s F(3.0, 4.11)=2.23, p=0.2240; IG2 Welch’s F(2.0, 9.30)=0.60, p=0.5709).

**Figure 4.**
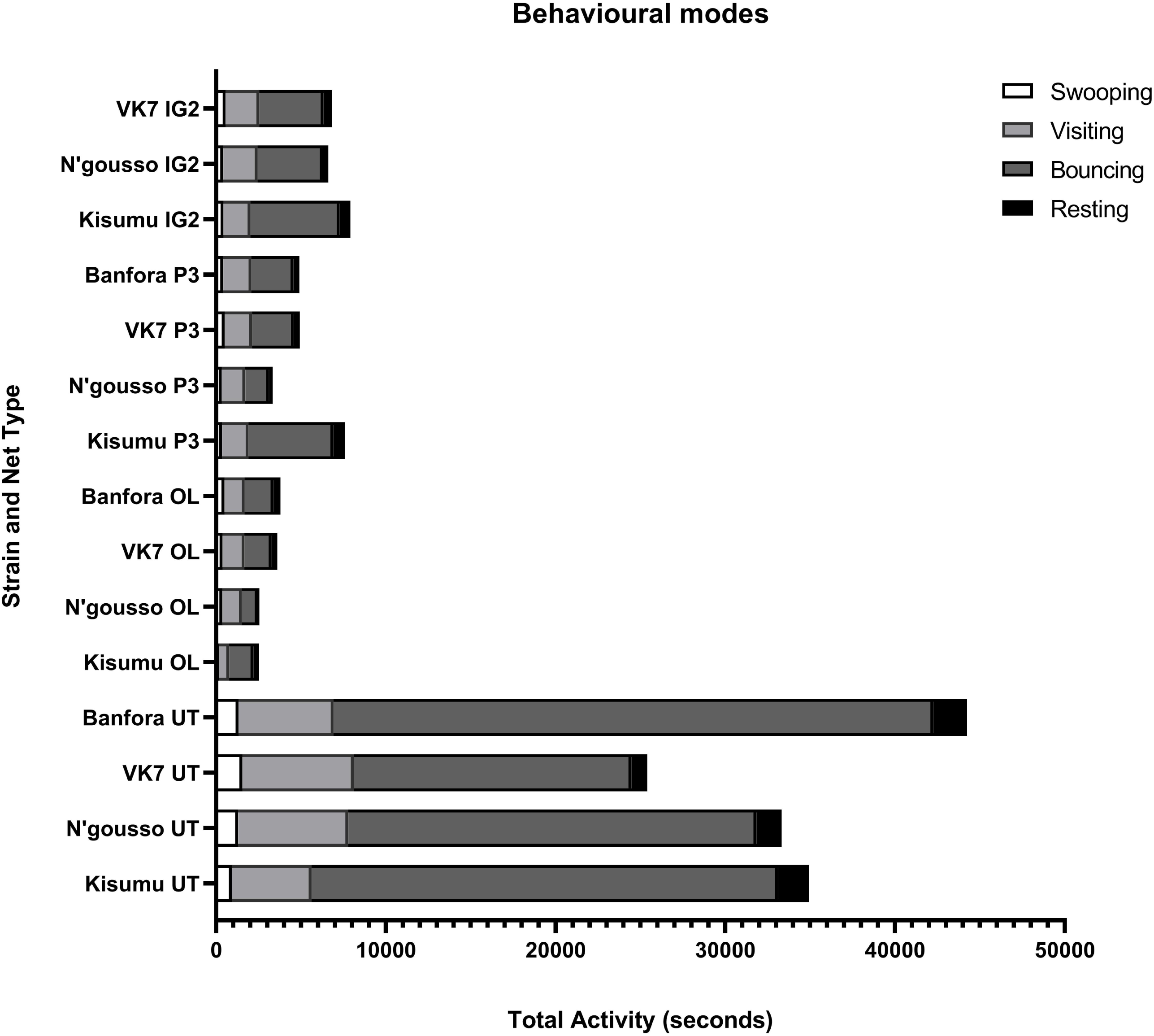
Behaviour of *Anopheles gambiae* at human baited bed nets. Mean total activity time of *Anopheles gambiae* recorded for each behavioural mode over two-hour recording period. As multiple mosquitoes were active simultaneously in the field of view, the total activity time could exceed the total recording time of 2 hours (7,200 seconds).

Breaking down total mosquito activity to look at time spent in each of the four distinct behavioural modes, revealed that both susceptible and resistant mosquitoes always spent more time swooping, visiting, bouncing and resting at an UT net than at any of the three ITNs (Additional Table 4; he one exception to this was comparing VK7 on UT and IG2, where there was no difference in total time spent resting (VK7 UT v IG2 p=0.1591)). However, there were no significant differences in the proportionate amounts of time spent swooping, visiting, bouncing, or resting between different ITNs (Additional Table 5).

Results comparing total activity changes on each net between strains for the four behavioural modes, showed that there was no difference in swooping activity between any strains on any nets, bar VK7 showing more activity than Kisumu around an UT net (UT Kisumu v VK7 p=0.0010). Analysis of total visiting time showed that N’gousso and VK7 spent more time in this behavioural mode than Kisumu when an UT net was present (UT Kisumu v N’gousso p=0.0352, Kisumu v VK7 p=0.0248), but there were no differences when comparing between any other nets. Banfora spent significantly more time bouncing on UT net than all other strains (UT Kisumu v Banfora p=0.0014, N’gousso v Banfora p<0.0001, VK7 v Banfora p<0.0001), and both susceptible strains spent more time bouncing than resistant VK7 (Kisumu v VK7 p<0.0001 N’gousso v VK7 p=0.0032). There was no difference in time spent bouncing between any strains on any of the ITNs. Kisumu and Banfora spent more time resting on an UT net than VK7 (UT Kisumu v VK7 p=0.0004, VK7 v Banfora p=0.0001), but there were no other significant differences in total time spent resting with an UT net or any of the ITNs (Additional Table 6).

### Quantifying number and duration of net contact

#### Contact number

All strains showed significantly greater mean total number of contacts with the UT net than with any of the ITNs (Additional Table 7). There were significant differences in the mean number of contacts with an UT net between some strains: Banfora had significantly more contact with the UT net than N’gousso and VK7, while Kisumu and N’gousso had more contact than VK7. Within strain comparisons showed there was no significant difference in the number of contacts made with any of the ITNs (Additional Table 8). There was also no difference in the number of contacts made between any of the strains on any of the ITNs (Additional Table 9) (Figure 5, panel A).

**Figure 5.**
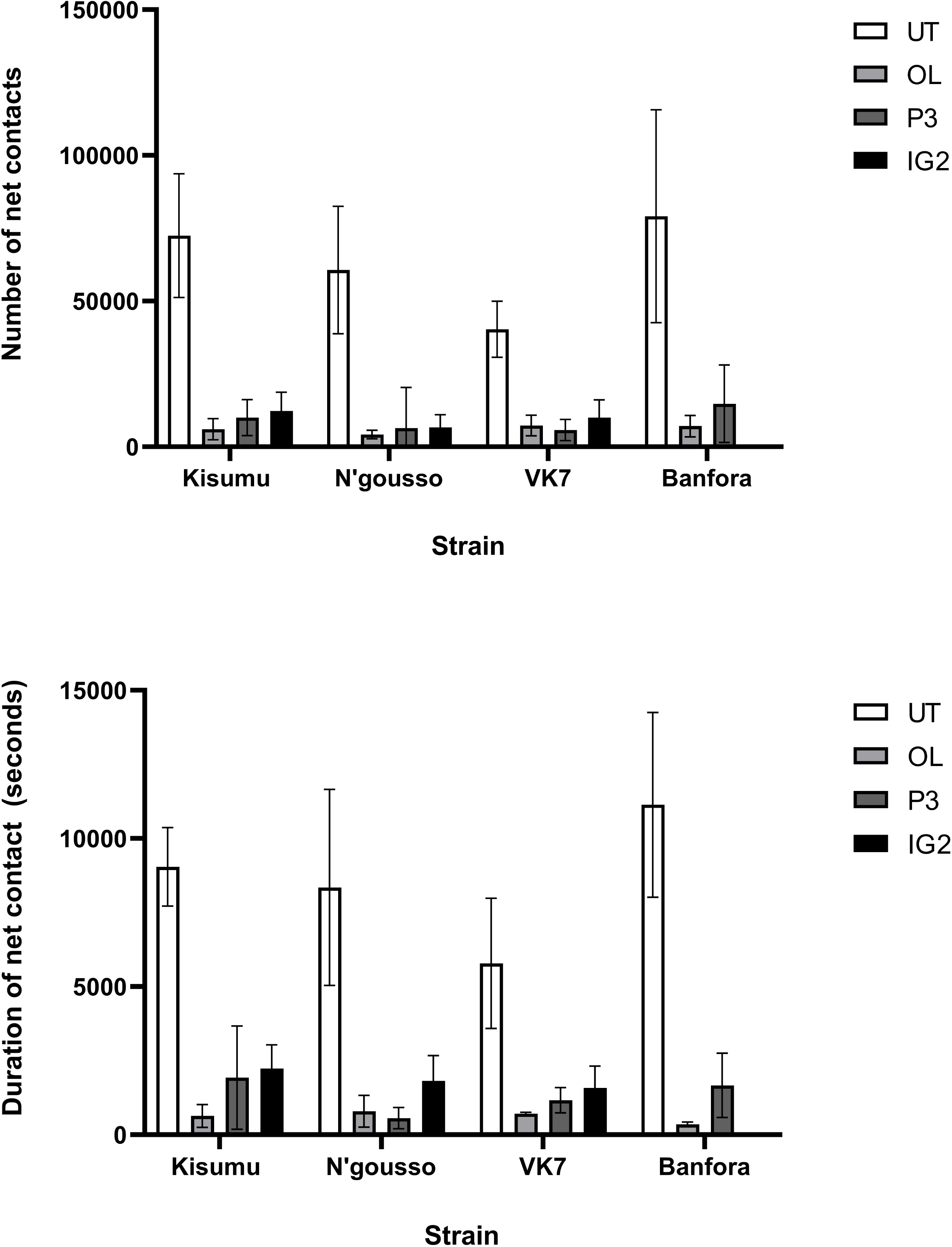
Mean total number of net contacts (A) and mean total duration net contact (B) with 95% Confidence Intervals for susceptible (Kisumu and N’gousso) and resistant (VK7 and Banfora) *Anopheles gambiae* strains on untreated net (UT), Olyset Net (OL), PermaNet 3.0 (P3) and Interceptor G2 (IG2).

#### Contact duration

Both susceptible and resistant mosquitoes spent significantly more time in contact with the UT net than any of the ITNs. Kisumu spent significantly more time in contact with IG2 than OL, but there were no other differences between nets (Additional Table 10). Between strain comparisons showed that Banfora spent significantly more time on UT net than all other strains, and both susceptible strains had longer contact duration than VK7. There was no significant difference in net contact duration for any strain combinations on treated nets (Additional Table 11) (Figure 5, panel B).

We calculated that during the 120-minute recording period each mosquito had between 285.62 seconds and 1041.79 seconds of contact with the UT net. There were no significant differences in the minimum and maximum time that susceptible and resistant mosquitoes spent on any of the three ITNs (OL: susceptible strains between 7.58 seconds and 101.39 seconds, resistant strains between 3.39 seconds and 255.53 seconds; P3: susceptible strains between 40.30 seconds to 241.77 seconds, resistant strains 33.35 seconds to 273.47 seconds; IG2: susceptible strains between 40.45 seconds and 403.39 seconds, resistant strain between 34.44 seconds and 378.73 seconds). The only notable differences we observed were that the minimum time that one Kisumu mosquito could have spent on OL was significantly lower than IG2 (p=0.0344), and the maximum time that N’gousso spent on IG2 was longer than on OL (p=0.0243) (Table 4).

**Table 4.** Minimum and maximum individual mosquito net contact duration (seconds) for entire 120minute recording.

| *Treatment* | *Strain* | *Minimum contact duration (s)* | *Maximum contact duration (s)* |
| --- | --- | --- | --- |
| **Untreated** | Kisumu | 301.45 | 952.28 |
|  | N'gousso | 398.72 | 962.83 |
|  | VK7 | 285.62 | 714.06 |
|  | Banfora | 542.17 | 1041.79 |
| **Olyset** | Kisumu | 7.58 | 101.39 |
|  | N'gousso | 9.96 | 64.28 |
|  | VK7 | 18.7 | 77.93 |
|  | Banfora | 3.39 | 255.53 |
| **P3** | Kisumu | 40.3 | 241.77 |
|  | VK7 | 33.35 | 273.47 |
|  | Banfora | 46.65 | 323.24 |
| **IG2** | Kisumu | 52.44 | 403.39 |
|  | N'gousso | 40.45 | 341.07 |
|  | VK7 | 34.44 | 378.73 |

### Net interactions in first 10minutes of assay

We investigated net contact in the first 10 minutes of the video tracking to examine if there was any suggestion of immediate repellent effects of the ITNs. While contact number and contact duration was lower at ITNs than UT nets, a higher percentage of overall contact duration occurred in the first 10 minutes of the assay on ITNs for the susceptible strains (Table 5). In the first 10 minutes, Kisumu spent significantly more time in contact with the ITNs than UT, and more time in contact with IG2 than OL or P3. Similarly, N’gousso had a higher percentage of contact time occurring in the first part of the assays when OL and IG2 were present, compared to the UT net. Again, N’gousso also had a longer contact duration on IG2 than OL. For resistant VK7, the highest initial 10-minute contact duration was observed on P3, whereas Banfora showed similar time spent across all three treatments.

**Table 5.**
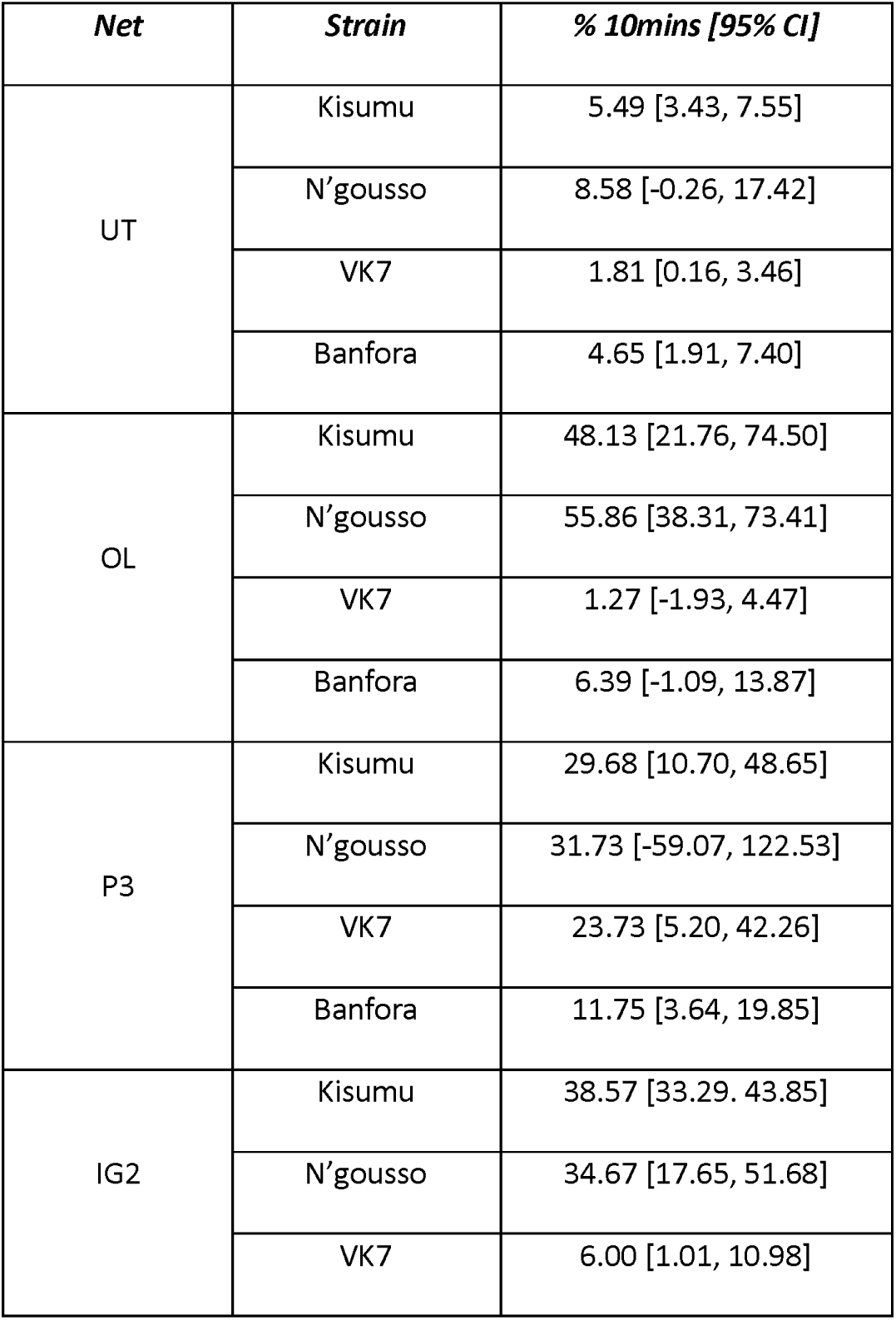
Percentage of overall contact duration occurring in the first 10minutes of the 2hour assay [95% Confidence Intervals].

| *Net* | *Strain* | *% 10mins [95% CI]* |
| --- | --- | --- |
| UT | Kisumu | 5.49 [3.43, 7.55] |
|  | N'gousso | 8.58 [-0.26, 17.42] |
|  | VK7 | 1.81 [0.16, 3.46] |
|  | Banfora | 4.65 [1.91, 7.40] |
| OL | Kisumu | 48.13 [21.76, 74.50] |
|  | N'gousso | 55.86 [38.31, 73.41] |
|  | VK7 | 1.27 [-1.93, 4.47] |
|  | Banfora | 6.39 [-1.09, 13.87] |
| P3 | Kisumu | 29.68 [10.70, 48.65] |
|  | N'gousso | 31.73 [-59.07, 122.53] |
|  | VK7 | 23.73 [5.20, 42.26] |
|  | Banfora | 11.75 [3.64, 19.85] |
| IG2 | Kisumu | 38.57 [33.29, 43.85] |
|  | N'gousso | 34.67 [17.65, 51.68] |
|  | VK7 | 6.00 [1.01, 10.98] |

Despite differences within strains on different nets, there were no differences observed between susceptible and resistant strains for 10-minute contact duration when an UT net or P3 was present. There was, however, a difference with OL, as both susceptible strains had a higher percentage of their overall contact duration occurring in this first period than both resistant strains. Susceptible strains also spent considerably more time contacting IG2 than VK7 (Additional Table 12, 13, 14, 15).

### Location of activity at the bed net interface

The distribution of total activity was heavily focused on the roof of the bed net for all strains and all net treatments (>90% on UT, >85% OL, >72% P3 and >87% IG2) as described in previous studies on standard ITNS (Lynd & Mccall, 2013; Parker et al., 2017) (Table 6). There was no significant difference in the percentage of contact occurring on the roof of the net for any strain or net combinations.

**Table 6.**
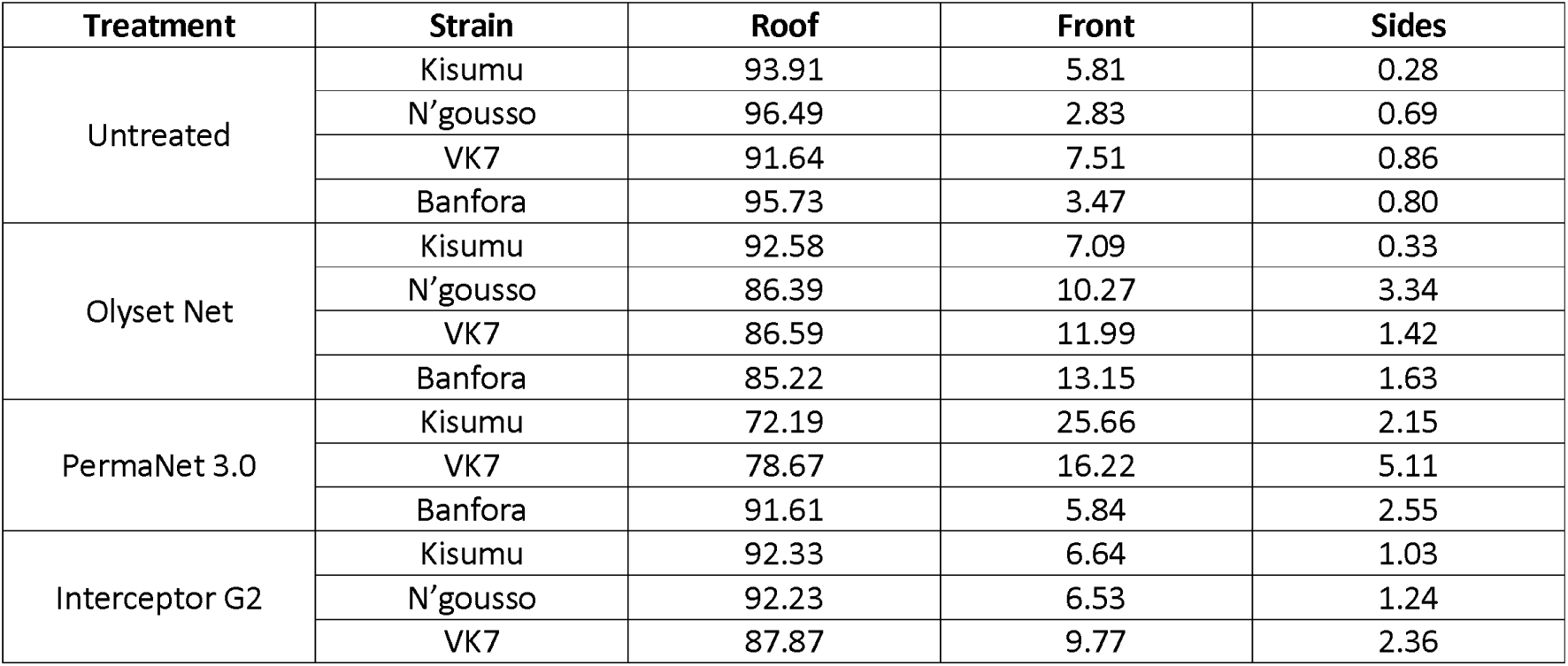
Percentage of overall contact across different regions of the bed net (%).

| Treatment | Strain | Roof | Front | Sides |
| --- | --- | --- | --- | --- |
| Untreated | Kisumu | 93.91 | 5.81 | 0.28 |
|  | N'gousso | 96.49 | 2.83 | 0.69 |
|  | VK7 | 91.64 | 7.51 | 0.86 |
|  | Banfora | 95.73 | 3.47 | 0.80 |
| Olyset Net | Kisumu | 92.58 | 7.09 | 0.33 |
|  | N'gousso | 86.39 | 10.27 | 3.34 |
|  | VK7 | 86.59 | 11.99 | 1.42 |
|  | Banfora | 85.22 | 13.15 | 1.63 |
| PermaNet 3.0 | Kisumu | 72.19 | 25.66 | 2.15 |
|  | VK7 | 78.67 | 16.22 | 5.11 |
|  | Banfora | 91.61 | 5.84 | 2.55 |
| Interceptor G2 | Kisumu | 92.33 | 6.64 | 1.03 |
|  | N'gousso | 92.23 | 6.53 | 1.24 |
|  | VK7 | 87.87 | 9.77 | 2.36 |

### Mosquito velocity during interaction with host within bed nets

Average speed of whole swooping tracks was analysed to assess changes in speed between strains around different bed nets. Only susceptible Kisumu showed any difference in flight speed around different net treatments, flying significantly faster around OL and IG2 than UT nets. Resistant strains did now show any difference in flight speed between different net types. Between strains, both resistant strains flew faster around an UT net than Kisumu and Banfora was significantly faster than Kisumu around P3. There was so difference in overall swooping speed between strains when OL or IG2 were present (Additional Table 16, 17).

### Mosquito interaction with the bed nets over time

We observed a steep decay in activity over the duration of the assay for susceptible strains with P3 and OL compared to UT net (Kisumu: UT v OL p=0.0023, UT v P3 p=0.0020). Kisumu also showed a dramatic decrease in activity in the presence of IG2 (UT v IG2 p<0.0001), which was not replicated in N’gousso activity decay around the same net. Resistant strains showed a less dramatic decay in activity when P3 and OL present, however decay was still more pronounced than with UT (VK7 UT v OL p=0.0128, UT v P3 p=0.0010), and there was no significant activity decay when VK7 was exposed to IG2. All strains exhibited no activity decay in the presence of an UT net (Figure 6) (Additional Table 18, 19).

**Figure 6.**
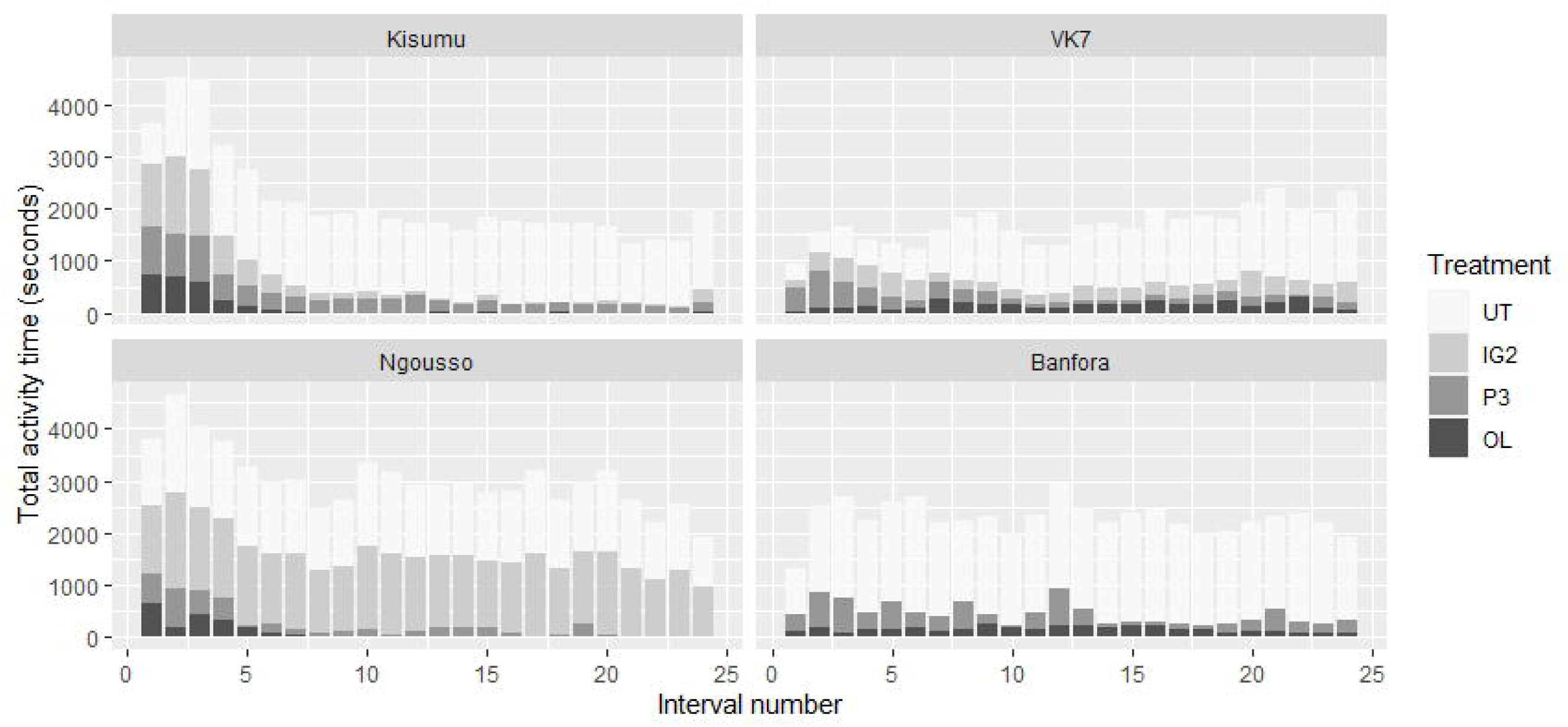
Rates of *Anopheles gambiae* activity across all four behavioural modes combined, throughout 120minute recording test period. Total activity is shown for untreated net (UT), Olyset Net (OL), PermaNet 3.0 (P3) and Interceptor G2 (IG2) for Kisumu, N’gousso, VK7 and Banfora.

### Sub-lethal pipeline – wing size and willingness to feed

Wing size was measured as a proxy for mosquito body size. There was a negative correlation between wing size and blood-feeding inhibition, with smaller mosquitoes less likely to survive and accept a bloodmeal. However, there was no significant interaction between wing size and strain (p=0.9447), suggesting that the relationship between wing size and blood feeding success was the same for all strains.

The majority of susceptible mosquitoes exposed to the three ITNs were either knocked-down or dead and hence unable to blood feed. OL reduced resistant strain feeding by up to 83% (VK7 71% [95% CI 62, 80], Banfora 83% [95% CI, 76, 91]), P3 reduced VK7 feeding by 97% [95% CI 94, 99], whereas IG2 had a smaller effect, reducing VK7 blood feeding success by 41% [95% CI 31, 51] (Figure 7). Between 14% and 70% of mosquitoes were unable to blood feed after exposure to UT net.

**Figure 7.**
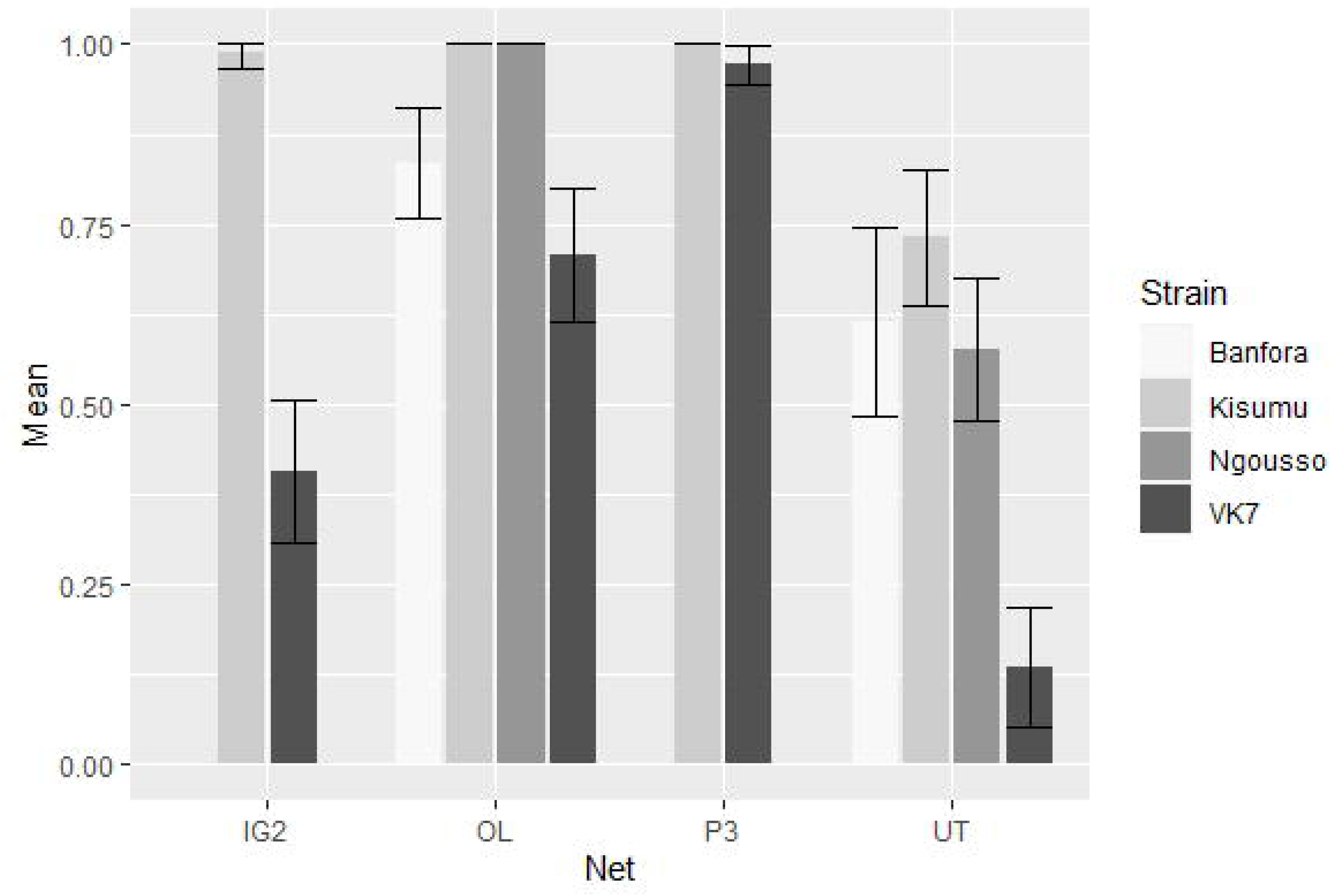
Predicted mean reduction in blood-feeding success of four strains after exposure to untreated (UT), Olyset Net (OL), PermaNet 3.0 (P3) and Interceptor G2 (IG2) (95% CI).

## Discussion

These results provide a first in-depth description of the behaviour of susceptible and resistant Anopheles gambiae strains around next-generation bed nets and the impact of these new nets on them. As insecticide resistance continues to be a growing threat to the success of African vector control programmes, there is an urgent need for safe novel treatments suitable for use on ITNs. The first of the next-generation nets using these treatments are now being evaluated in field trials (Mosha et al., 2022; Tungu et al., 2021) and deployed at scale in pilot studies in several countries (IVCC, 2020). Determining how mosquitoes interact with the nets, and the consequences of net contact for mosquitoes, will aid in interpretation of the results of clinical trials, and extrapolation to alternative settings with different mosquito populations.

OL, P3 and IG2 all killed more than 90% of susceptible mosquitoes 24 hours after a 2-hour exposure, but this effect was not seen with resistant mosquitoes where only 20.4% of VK7 and 45.4% of Banfora on OL, 71.4% of VK7 and 72.4% of Banfora on P3, and 15.9% of VK7 on IG2 (the Banfora strain was not tested on this net) were dead at 24hours. Total mosquito activity was higher around an UT net than all ITNs, which is comparable with results obtained in previous studies (Parker et al., 2015). Interestingly, there was no difference in total activity observed between susceptible and resistant strains around any of the ITNs tested, the number and duration times of net contact was also similar for all strains. Net contact was focussed predominantly on the roof for all types of bed net and did not change throughout the assay (Parker et al., 2015; Sutcliffe & Yin, 2014). Through comparing the difference in the first and last 10 minutes of recording activity, we observed a steep decay in activity for both susceptible strains when P3 and OL were present, but only a decrease in activity around IG2 for susceptible Kisumu. Resistant strains showed a less dramatic decay in activity when P3 and OL present, however decay was still more pronounced than with UT. The activity decay in susceptible strains most likely reflects that mosquitoes are being knocked down and killed by the active-ingredients, however, the lack of decay observed with resistant strains is surprising, particularly for dual-treated nets.

The behaviour of the strains as measured by tracking was remarkably similar for all the strains tested with no significant differences observed in the number of contacts, or the duration of time spent contacting ITNs between susceptible and resistant mosquito strains. We did not observe evidence of a repellent effect on susceptible mosquitoes for any ITN as a higher percentage of overall contact duration occurred during the first 10 minutes of the assay on all ITNs compared to untreated net.

The low mortality results in resistant strains from our study do not match those from recent experimental hut studies reporting promising results with the Interceptor G2 net (Bayili et al., 2017; Camara et al., 2018; N’Guessan et al., 2016; Tungu et al., 2021) where mortality in huts with IG2 was significantly higher than for standard pyrethroid only ITNs in all settings. A recent clinical trial by Mosha et al., (2022) reported after two years IG2 provided significantly better protection from malaria than an alpha-cypermethrin only ITN in areas where mosquito populations are resistant to pyrethroids (Mosha et al., 2022). Nevertheless, when tested in a laboratory under standard conditions, the results from ours and other studies are not dissimilar, with low mortalities of insecticide resistant mosquitoes at both 24hours and 72hours post IG2 exposure. We recorded 25.6% mortality at 72hours in the resistant VK7 strains and others have reported ~5-26% mortality using a 3minute WHO cone assay and a wider range of between ~18% - 100% after a 30minute exposure in a WHO tube assay(Camara et al., 2018; N’Guessan et al., 2016). The reasons for differences in performance of IG2 under laboratory and field settings are unclear but differences in the mosquito population assessed may be important. Unpublished data from multiple experimental hut studies in southwest Burkina Faso (the region of origin of the VK7 and Banfora strains used in the current study) show relatively poor performance of IG2 nets compared to data from other settings (Sanou, A, Sagnon N, Guelbeogo M).

Moreover, the complete entomological mode of action of chlorfenapyr has not yet been determined and reproducing in the level of mortality seen in hut trials in laboratory assays has proven challenging. This severely limits our ability to apply lab tests, including video tracking, in the evaluation of products containing this insecticide.

Mosquitoes were given the opportunity to blood feed one-hour post-assay and we observed a reduction in blood feeding success with resistant strains after exposure to all ITNs. Despite lower mortality with the pyrethroid only Olyset Net and next-generation Interceptor G2, blood feeding success in resistant strains was reduced by up to 83% and 41% respectively. A reduction in blood-feeding after insecticide exposure was also found by Barreaux et al., (2022) who reported that that after forced exposure to ITNs the blood feeding success of highly insecticide resistant An. gambiae strains was reduced. The authors suggest that this was not a result of mosquitoes avoiding the net or being repelled by it, but instead because contact with insecticides reduced feeding capacity.

As previously observed (Parker et al., 2017), both susceptible and resistant strains showed a much higher level of overall activity when an UT net was present, with activity levels reducing dramatically in the presence of all tested ITNs. This reduction in activity was observed for all strains with no significant differences in total activity level between any of the strain and ITN comparisons. This suggests that even if this measurable reduction in activity is attributable to the pyrethroid component on the net the additional AI.s do not alter it. Moreover, the novel chemistries do not affect mosquitoes of differing resistance status differently. One result to note, is that despite the low mortality rate of VK7 when exposed to IG2, the time spent resting on this ITN was similar to that of when an UT net was present. One explanation for these results could be that there is a currently unknown interaction occurring between the two insecticides, which is reducing the efficacy of chlorfenapyr. We believe that this could be due to the pyrethroid suppressing chlorfenapyr activation by preferentially binding cytochrome p450s and hence delaying activation to the lethal metabolite tralopyril.

There are a few limitations to this study which are important to consider. While the environment in which the tracking assay data are collected reproduces as much as possible the conditions in the interior of a hut, there are important omissions and differences. Firstly, the (apparent) repellent properties of some nets that reduce initial eave entry cannot be measured here nor can the proportion of mosquitoes that leave the room after contacting the net. Hence all 25 mosquitoes must enter and remain in the room potentially delivering an overestimate of the lethality of the net being tested. Environmental conditions also remained static throughout the test whilst in reality air disturbances, and changes in temperature during the night may affect net contact.

It was not possible to determine individual mosquito contact, and total net contact was calculated based on the maximum number of mosquitoes seen simultaneously contacting the net in any one frame of the recording. Although this method provides a more realistic estimate of mosquito/ITN contact times than other standard bioassays, the measurement does not account for mosquitoes that make zero contact or that return to make multiple contacts with the net. This is especially important for the interpretation of sublethal results with contact duration varying between the individuals exposed. There is therefore a strong argument for collecting data to determine LD50 equivalents for duration of net contact, determined for each ITN. The video recordings in this study were limited to 2hours as the data files produced are extremely large (2-3Tb per camera, per recording), but recording mosquito behaviour for longer periods to assess any delayed effects on mosquito behaviour could prove important when evaluating impacts of nets with poorly understood AIs. Future studies would benefit from more replicates with multiple different resistant mosquito strains, to investigate the potential effect of different resistance mechanisms, an aspect of evaluating ITNs already supported by many (Lees et al., 2022).

Overall, these findings expand our knowledge of how mosquitoes interact with ITNs, particularly with regards to behaviour around new chemistries. These results indicate that the effects of a range of ITNs on mosquito behaviour is remarkably consistent with no major alterations in mosquito responses, particularly ITN contact resulting from exposure to the nets by strains of differing pyrethroid susceptibilities. It also appears that lower ITN contact is not the reason for observed lower mortality in resistant strains. Ongoing work in multiple field sites will continue to explore the effects of new ITNs on the behaviour of wild mosquito populations and will undoubtedly contribute to the body of work gathering as a foundation for understanding behavioural mechanisms of resistance.

## Author contributions

Conceptualisation: PJM, HR, GMF. Funding acquisition: PJM, HR, GMF. Hardware and software: DT, CET, VV. Data collection: KG, AG, ME, AnM. Data analysis: KG, AG, FM, AM. Writing, original draft: KG. Writing, review and editing: all authors.

## Conflicts of interest

Authors declare no conflicts of interest.

## Acknowledgements

We would like to thank all the volunteers who partook in this study.

## Funding

This research was funded by the Bill & Melinda Gates Foundation [under Grant Agreement Nos OPP1159078 and INV010445]. The findings and conclusions contained within are those of the authors and do not necessarily reflect positions or policies of the Bill & Melinda Gates Foundation.

## Availability of data and materials

The data sets used for the work in this article are available upon request from the authors.

## Additional material

**Additional Table 1.**
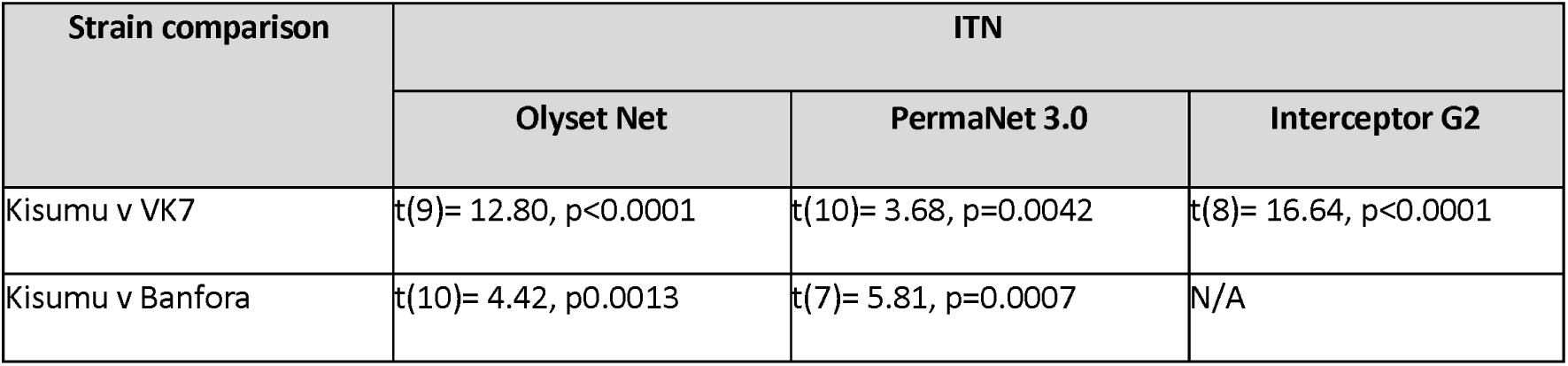
Mean 24hour mortality [95% CI]

**Additional Table 2.**
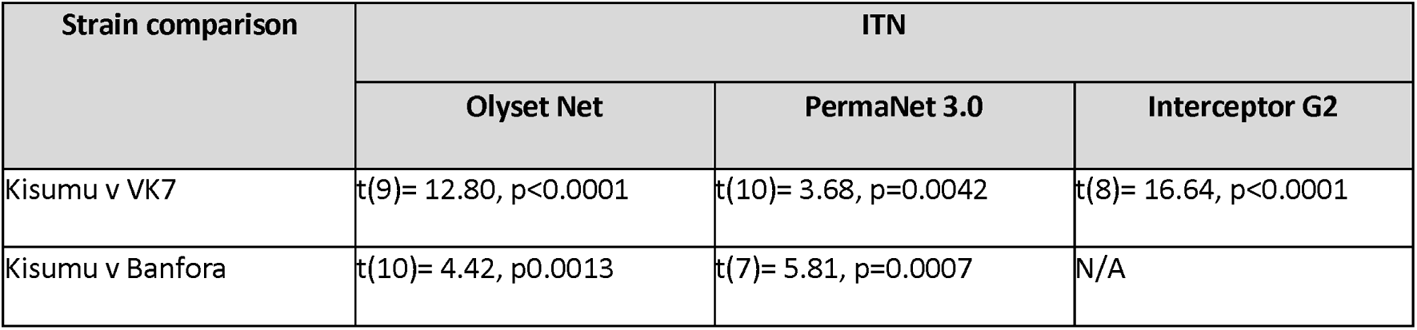
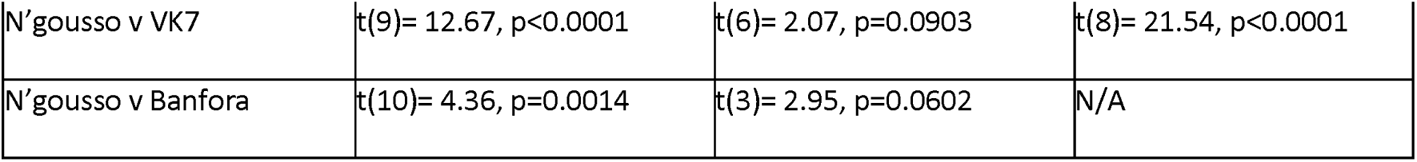
Mean 24hour mortality comparisons between three insecticide treated nets and four mosquito strains, two susceptible (Kisumu and N’gousso) and two resistant (VK7 and Banfora).

**Additional Table 3.**
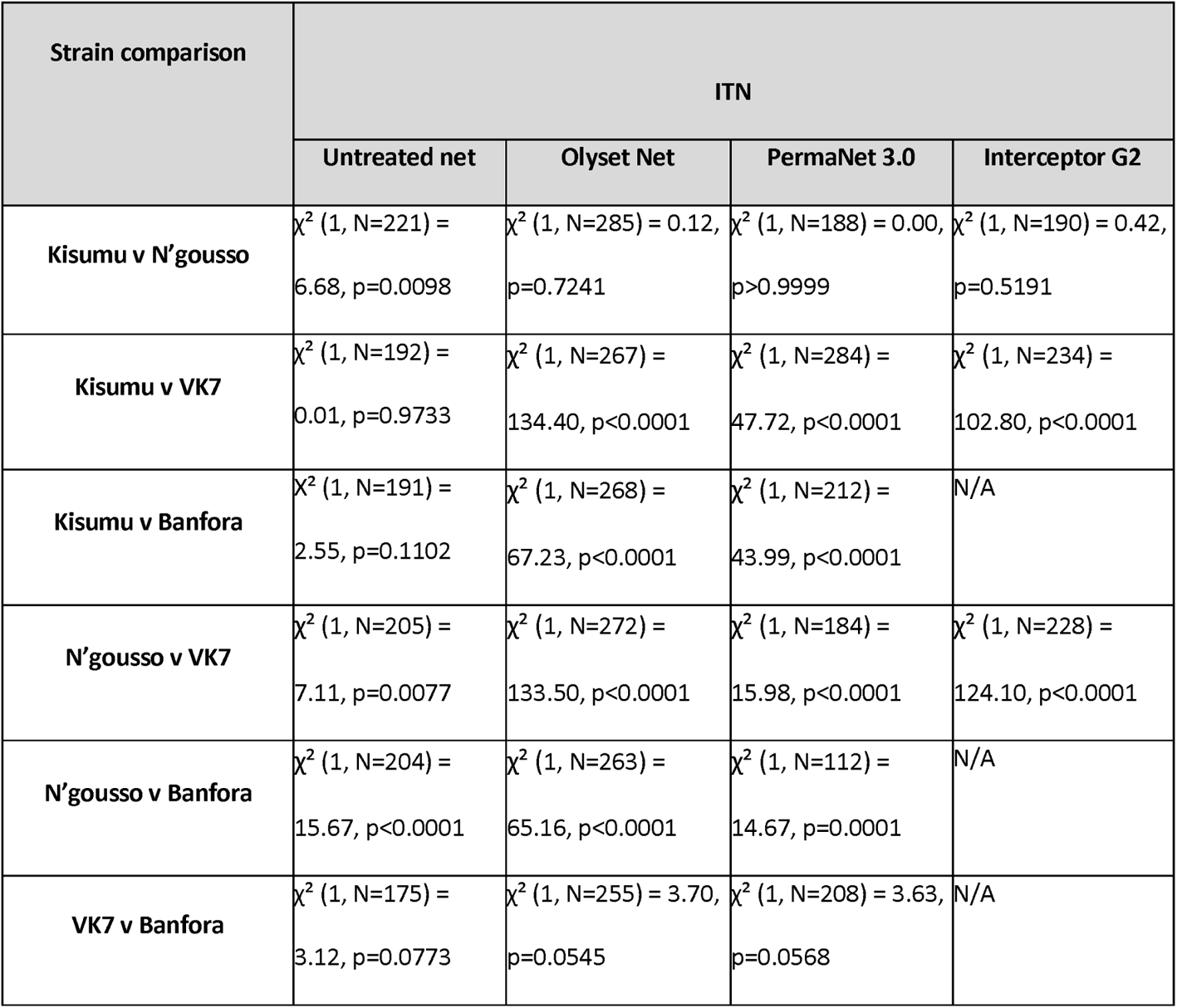
Comparison of median survival times of susceptible (Kisumu and N’gousso) and resistant (VK7 and Banfora) strains on four different net treatments.

| Strain comparison | ITN |  |  |  |
| --- | --- | --- | --- | --- |
|  | Untreated net | Olyset Net | PermaNet 3.0 | Interceptor G2 |
| **Kisumu v N'gousso** | $\chi^2$ (1, N=221) = 6.68, p=0.0098 | $\chi^2$ (1, N=285) = 0.12, p=0.7241 | $\chi^2$ (1, N=188) = 0.00, p>0.9999 | $\chi^2$ (1, N=190) = 0.42, p=0.5191 |
| **Kisumu v VK7** | $\chi^2$ (1, N=192) = 0.01, p=0.9733 | $\chi^2$ (1, N=267) = 134.40, p<0.0001 | $\chi^2$ (1, N=284) = 47.72, p<0.0001 | $\chi^2$ (1, N=234) = 102.80, p<0.0001 |
| **Kisumu v Banfora** | $\chi^2$ (1, N=191) = 2.55, p=0.1102 | $\chi^2$ (1, N=268) = 67.23, p<0.0001 | $\chi^2$ (1, N=212) = 43.99, p<0.0001 | N/A |
| **N'gousso v VK7** | $\chi^2$ (1, N=205) = 7.11, p=0.0077 | $\chi^2$ (1, N=272) = 133.50, p<0.0001 | $\chi^2$ (1, N=184) = 15.98, p<0.0001 | $\chi^2$ (1, N=228) = 124.10, p<0.0001 |
| **N'gousso v Banfora** | $\chi^2$ (1, N=204) = 15.67, p<0.0001 | $\chi^2$ (1, N=263) = 65.16, p<0.0001 | $\chi^2$ (1, N=112) = 14.67, p=0.0001 | N/A |
| **VK7 v Banfora** | $\chi^2$ (1, N=175) = 3.12, p=0.0773 | $\chi^2$ (1, N=255) = 3.70, p=0.0545 | $\chi^2$ (1, N=208) = 3.63, p=0.0568 | N/A |

**Additional Table 4.**
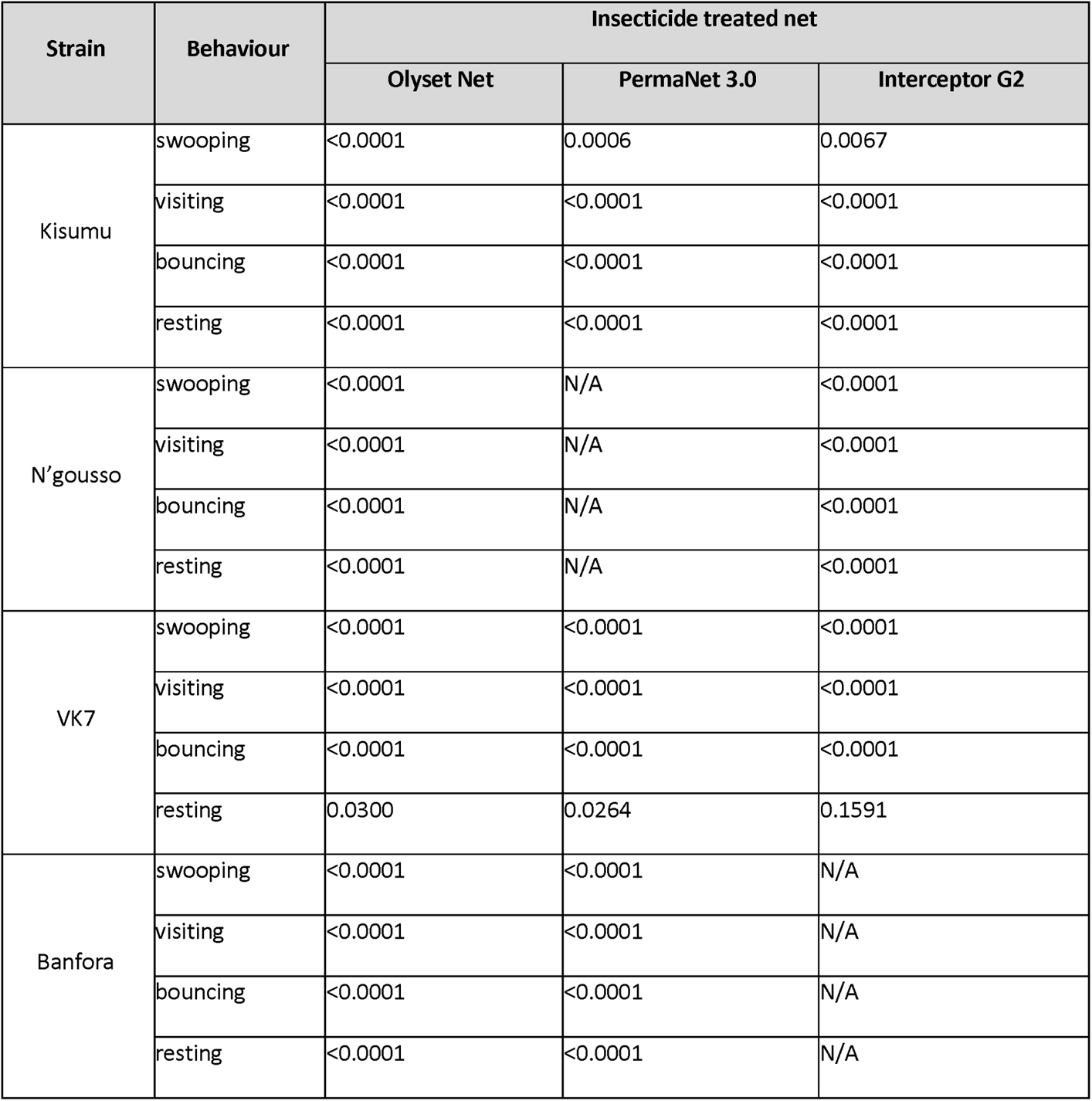
Statistically significant differences (p values) in total activity time split into four different behavioural modes (swooping, visiting, bouncing and resting), comparing untreated (UT) net to either Olyset Net (OL), PermaNet 3.0 (P3) or Interceptor G2 (IG2), for susceptible (Kisumu and N’gousso) and resistant (VK7 and Banfora) mosquitoes.

**Additional Table 5.**
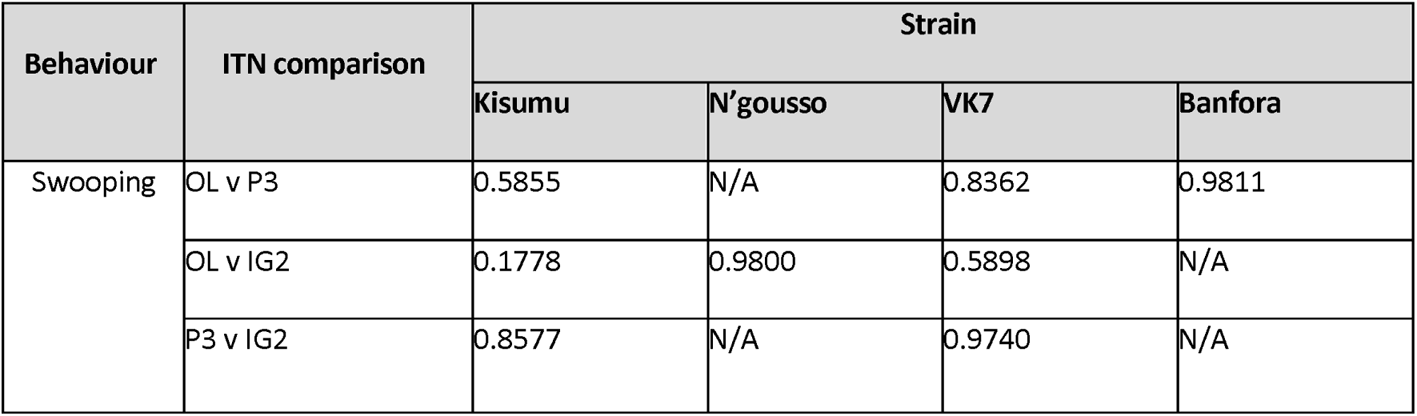
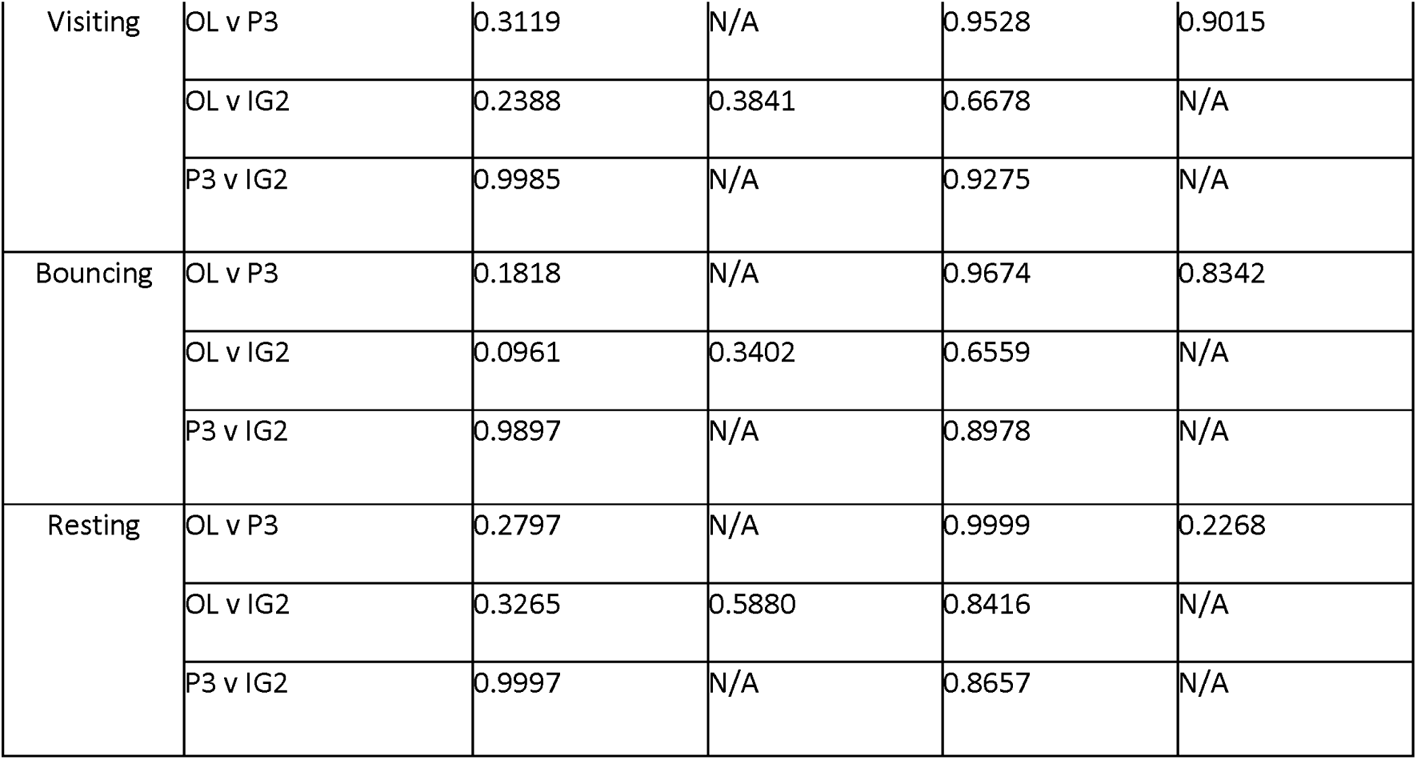
Within strain comparisons (p-value) of total activity time split into four different behavioural modes (swooping, visiting, bouncing and resting) between three ITNs (Olyset Net = OL, PermaNet 3.0 = P3, Interceptor G2 = IG2).

**Additional Table 6.**
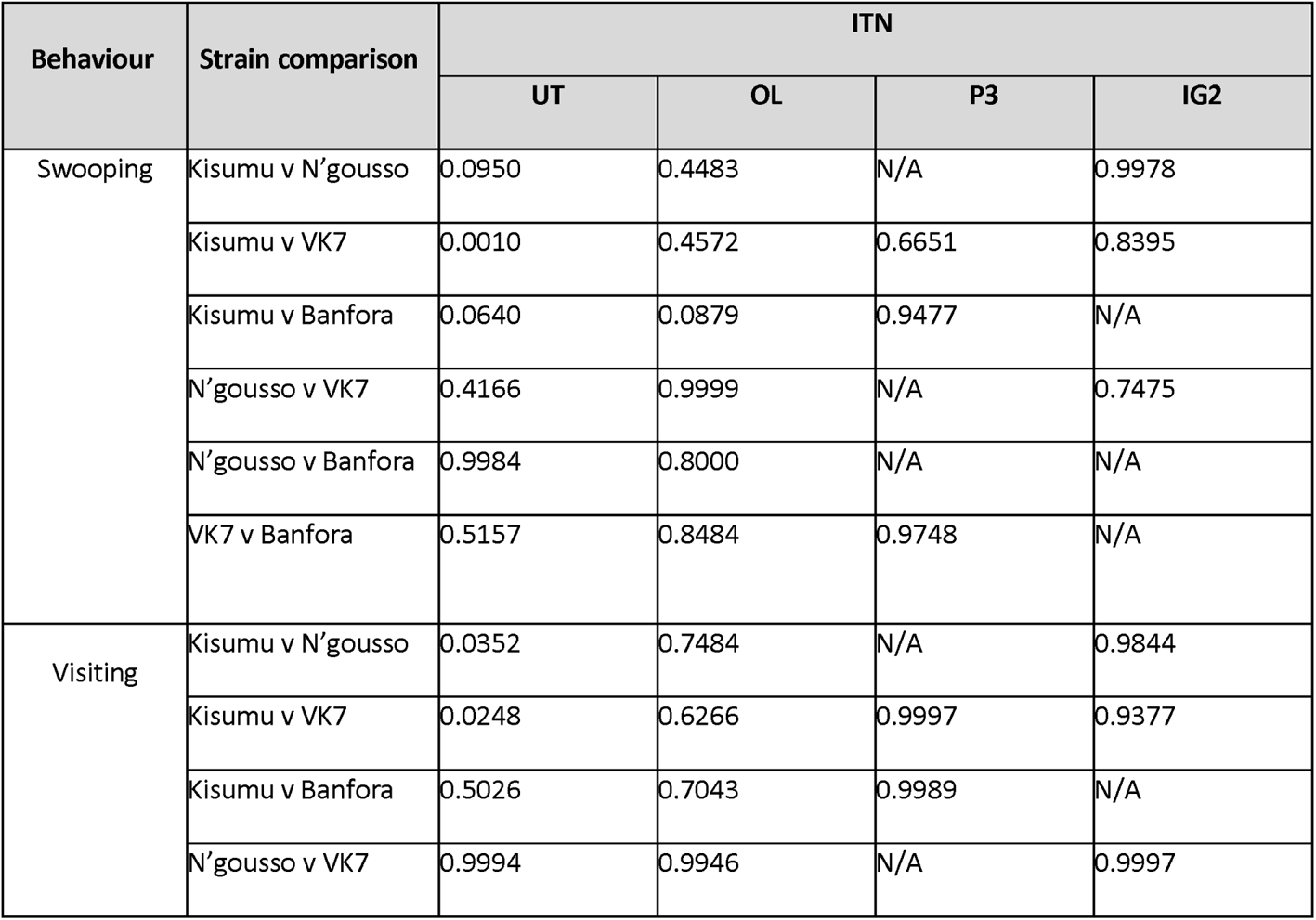
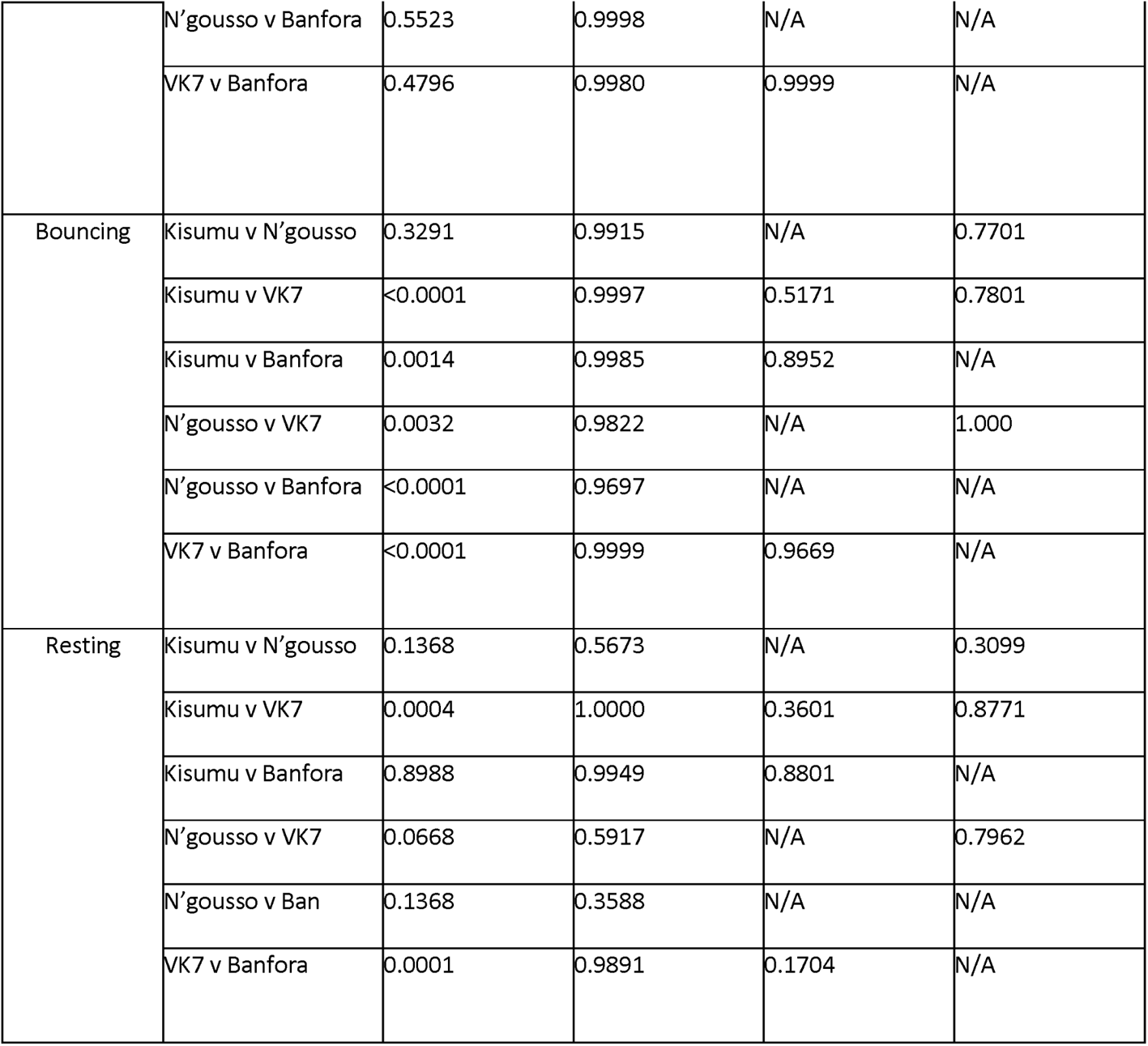
Within treatment comparisons (p-value) of total activity split into for behavioural modes (swooping, visiting, bouncing and resting) on four ITNs (Untreated net = UT, Olyset Net = OL, PermaNet 3.0 = P3, Interceptor G2 = IG2) between four mosquito strains

**Additional Table 7.**
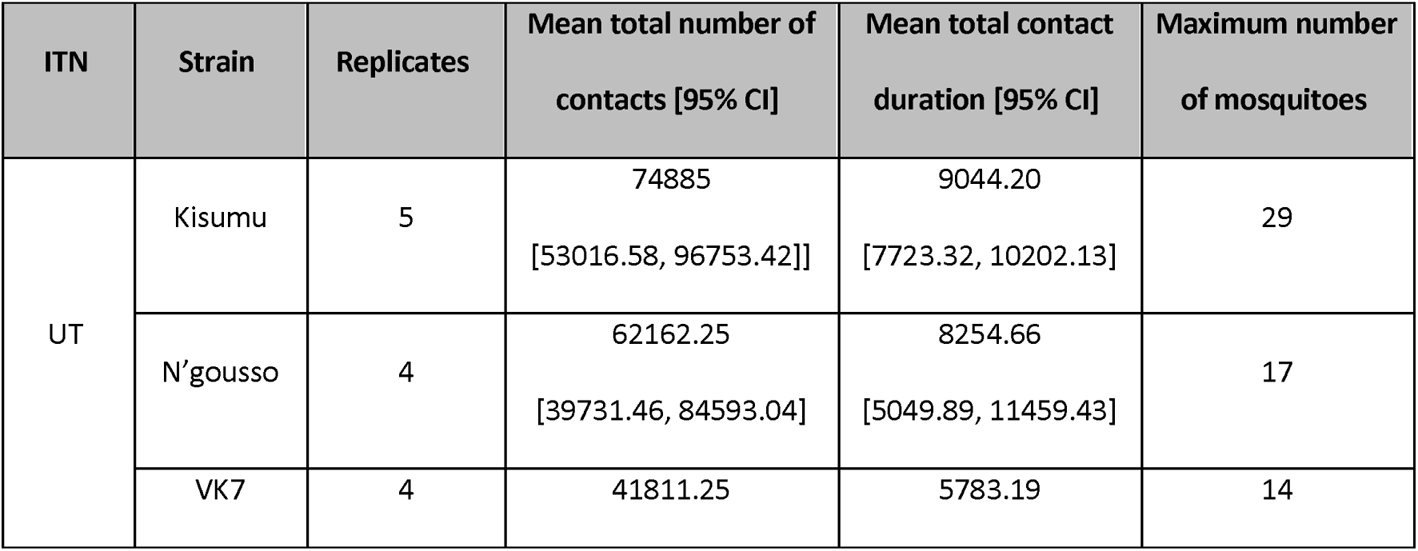
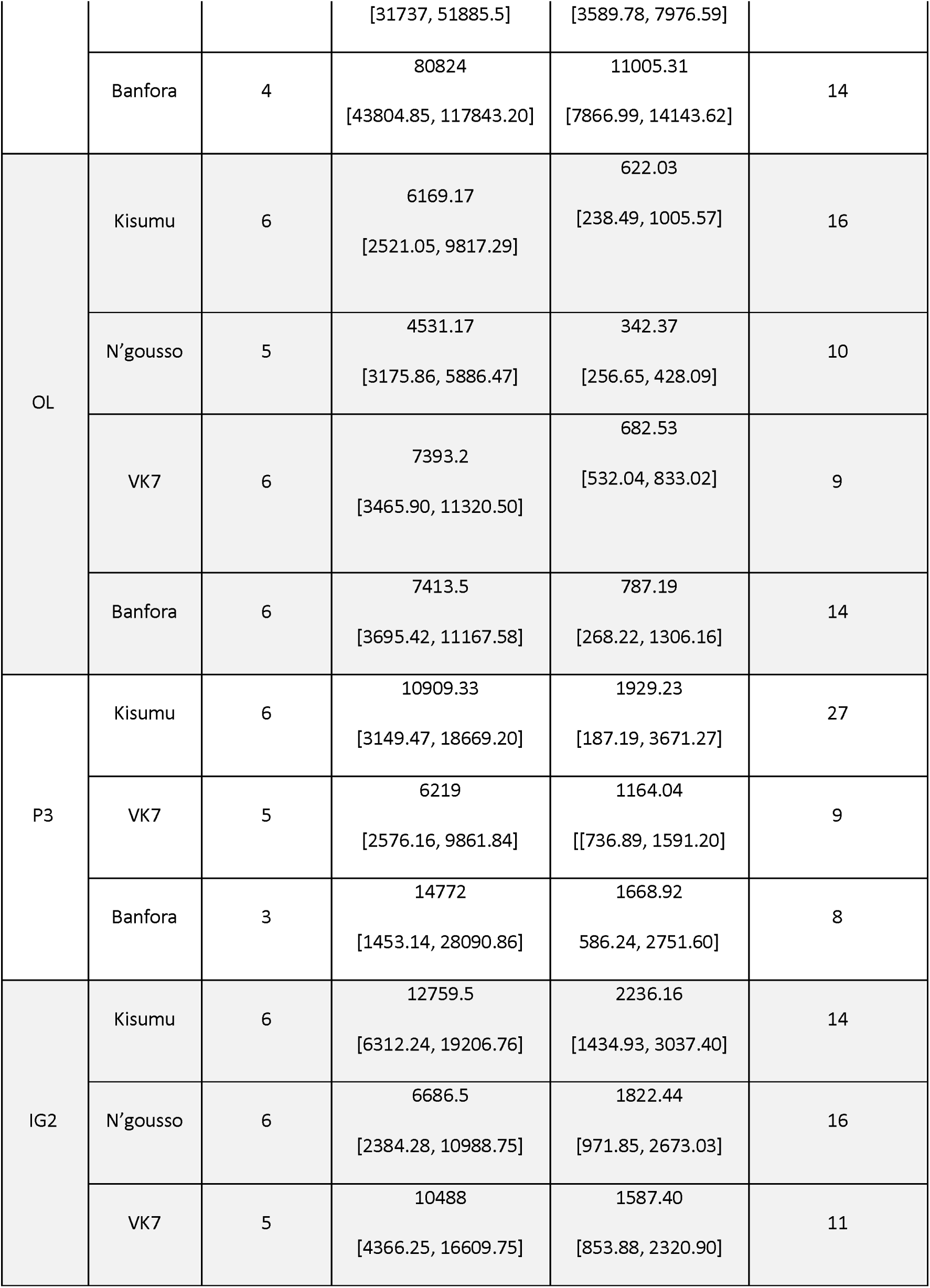
Mean total number of bed net contacts [95% CI], mean total contact duration [95% CI] and maximum number of mosquitoes seen in one frame of video recording.

**Additional Table 8.**
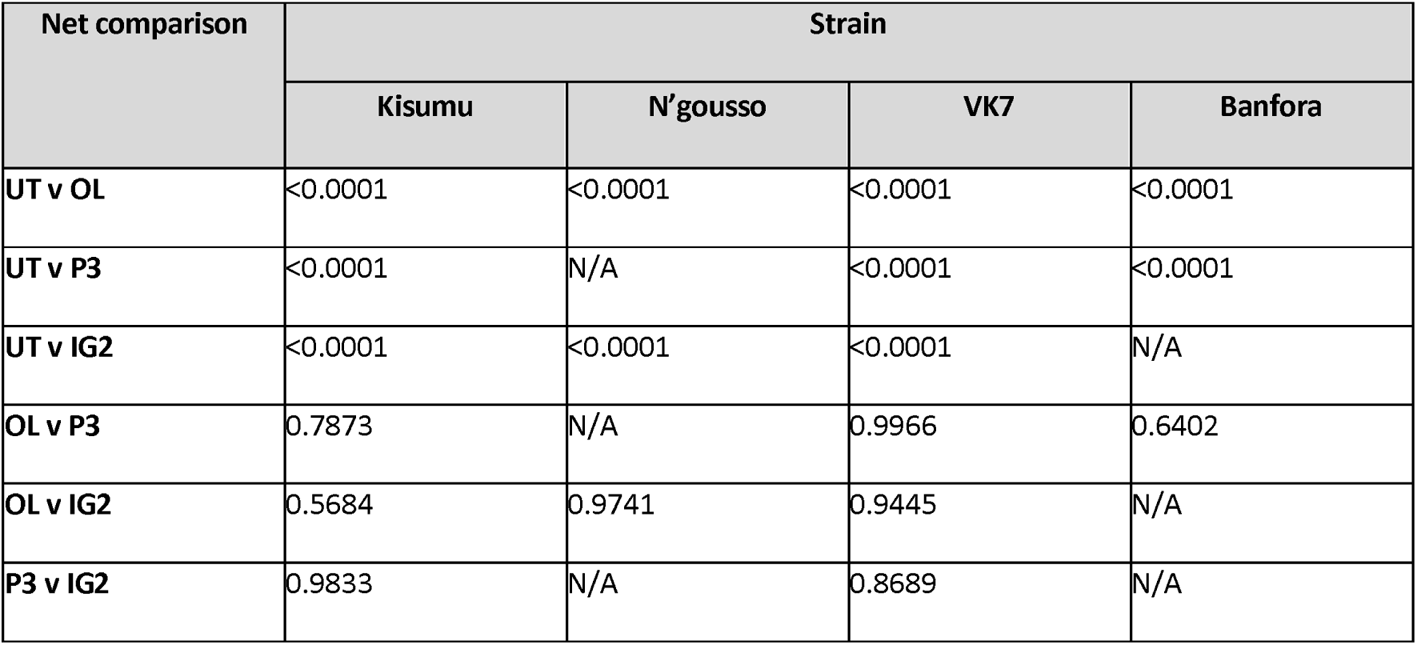
Within strain statistical comparisons (p value) of total number of net contacts for susceptible (Kisumu and N’gousso) and resistant (VK7 and Banfora) mosquitoes between four nets (UT = untreated, OL = Olyset Net, P3 = PermaNet 3.0, IG2 = Interceptor G2).

**Additional Table 9.**
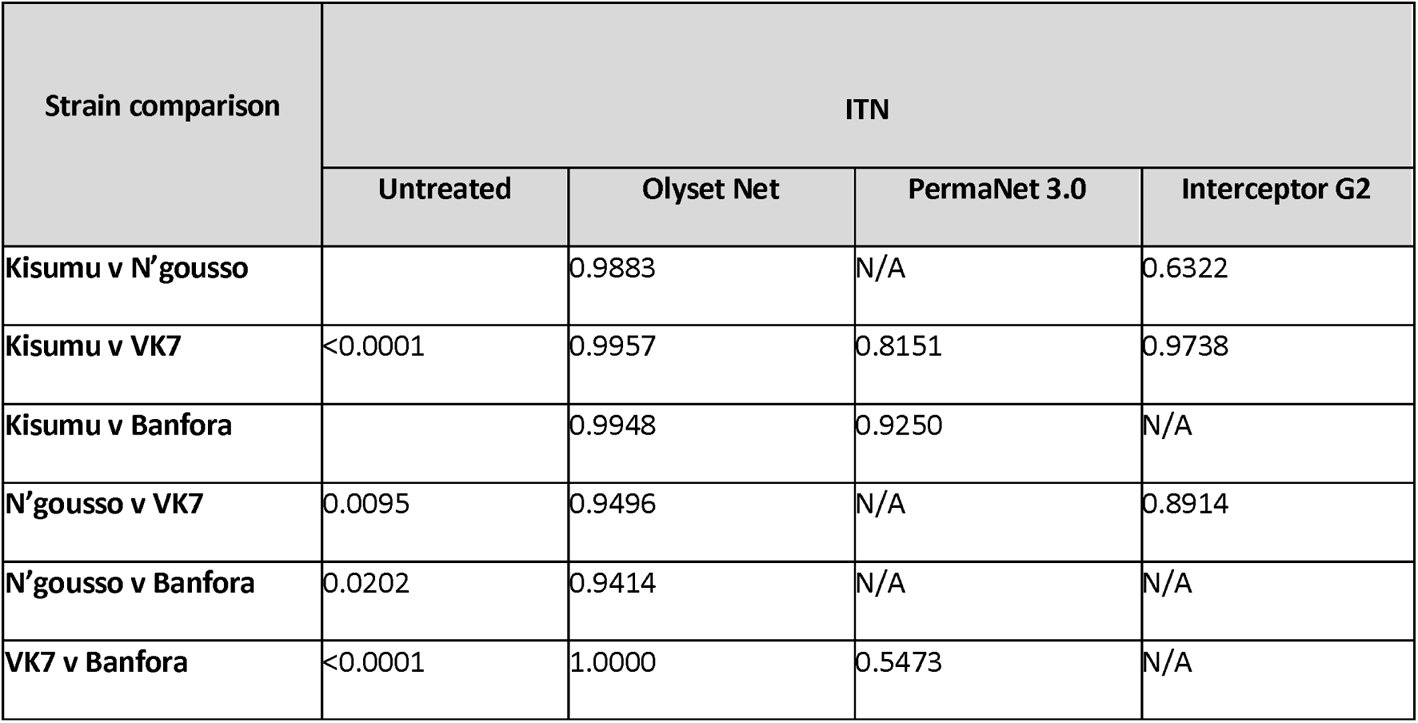
Within treatment statistical comparisons (p value) of total number of net contacts for four nets between four mosquito strains.

**Additional Table 10.**
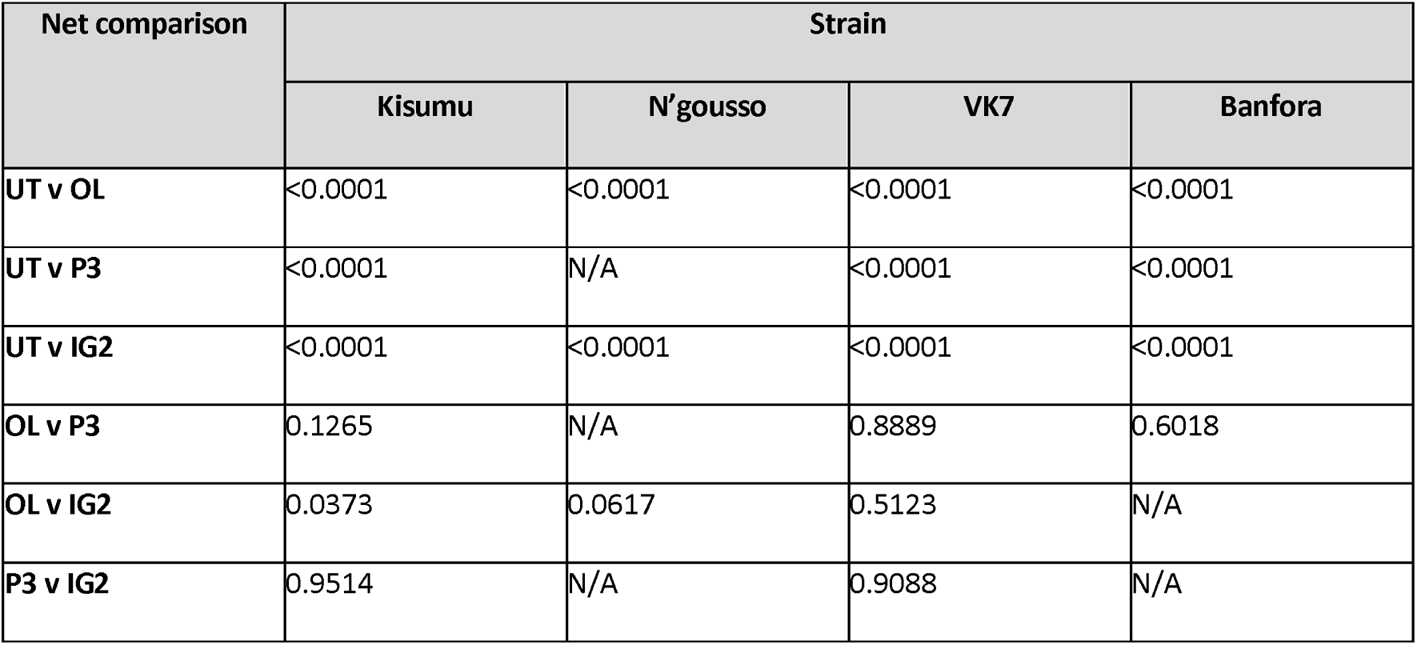
Within strain comparisons (p-value) of total duration of net contact for susceptible (Kisumu and N’gousso) and resistant (VK7 and Banfora) mosquitoes between three ITNs (OL = Olyset Net, P3 = PermaNet 3.0, IG2 = Interceptor G2).

**Additional Table 11.**
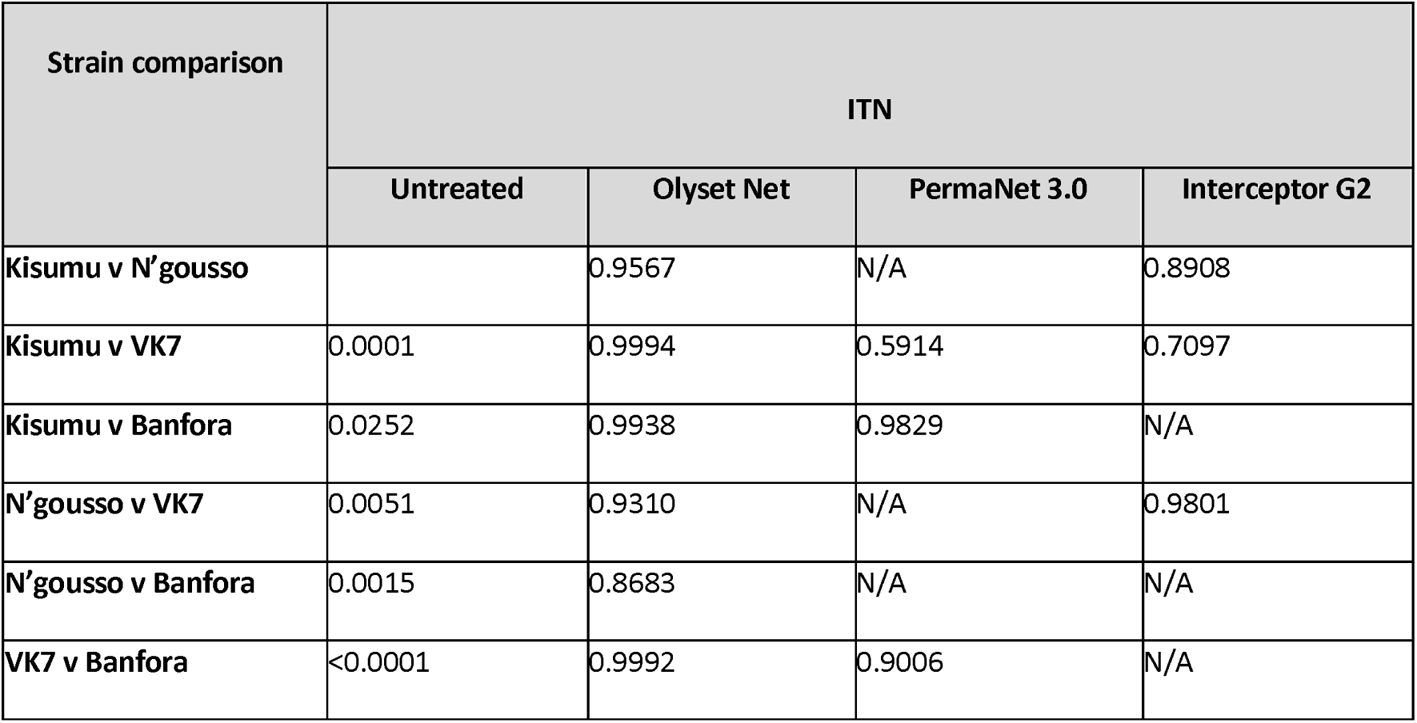
Within treatment comparison (p-value) of total net contact duration for three ITNs between four mosquito strains.

**Additional Table 12.**
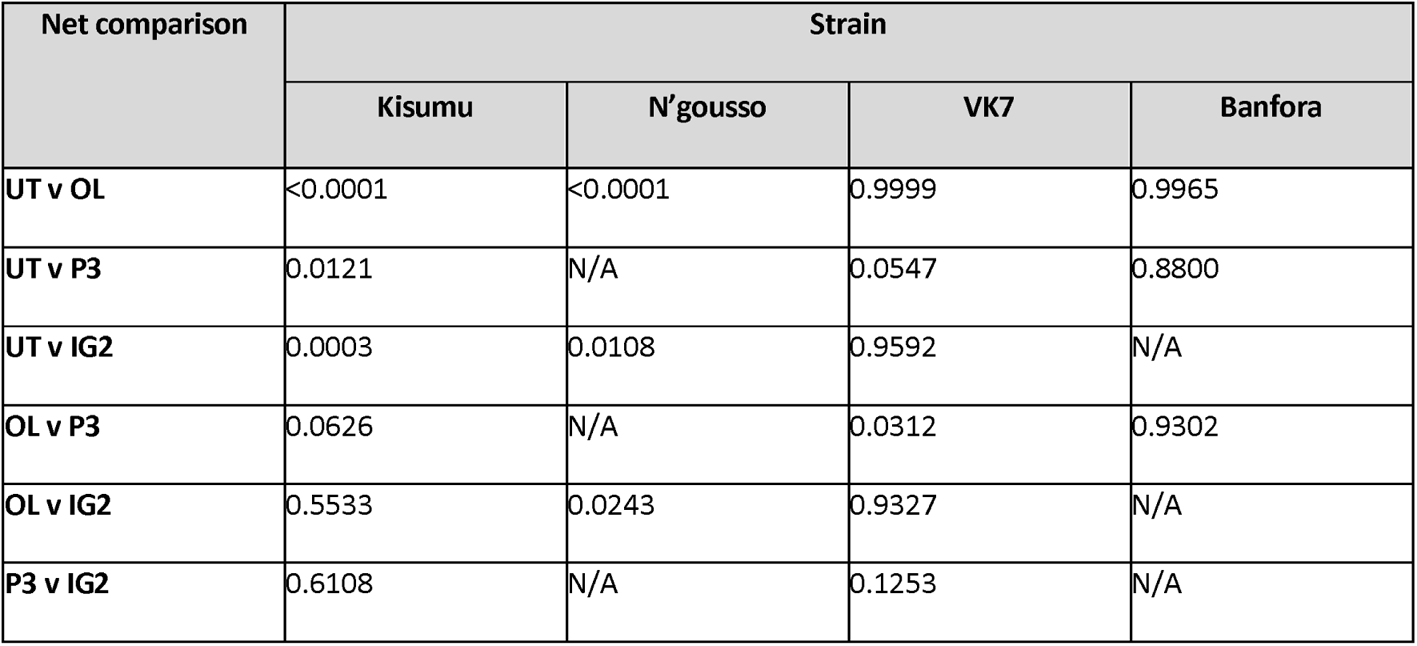
Percentage of contact duration in first the 10minutes of room scale tracking assay – within strain, between net differences.

**Additional Table 13.**
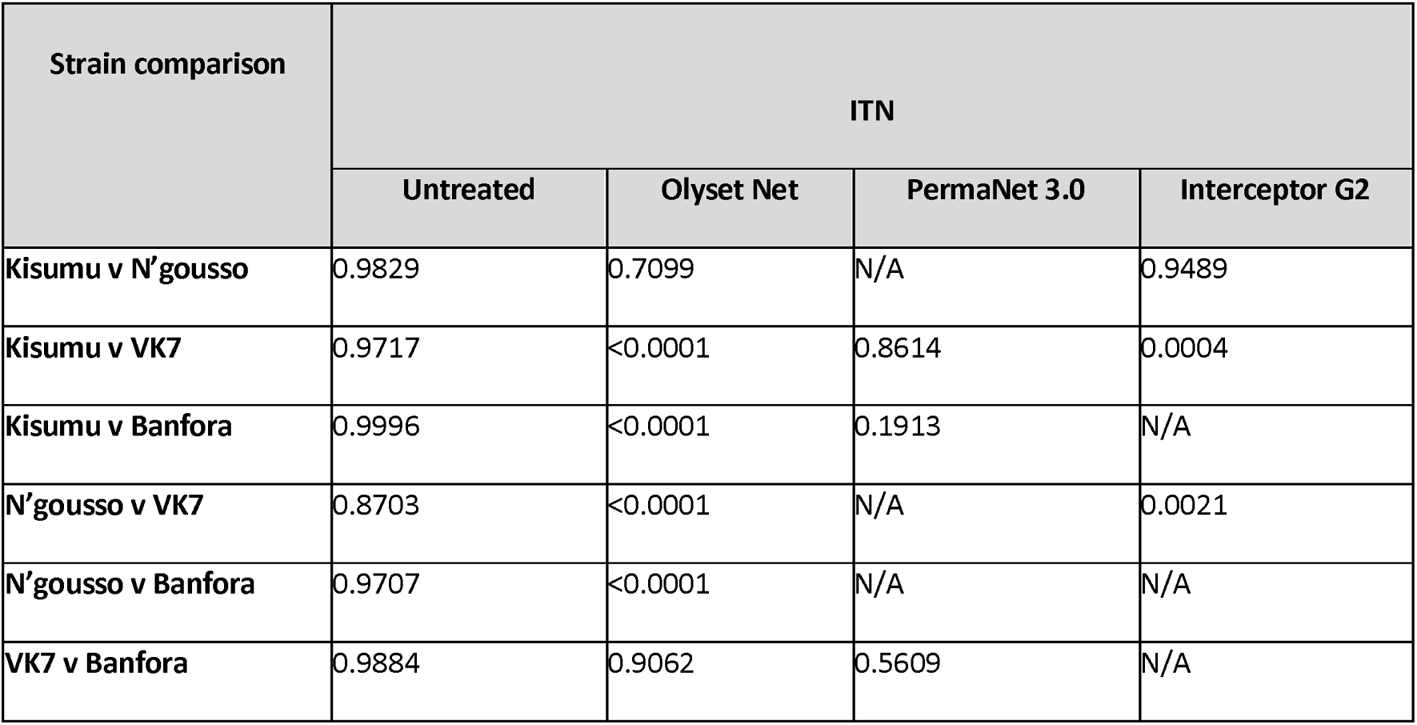
Percentage of contact duration in first 10mins of assay – within net, between strain differences

**Additional Table 14.**
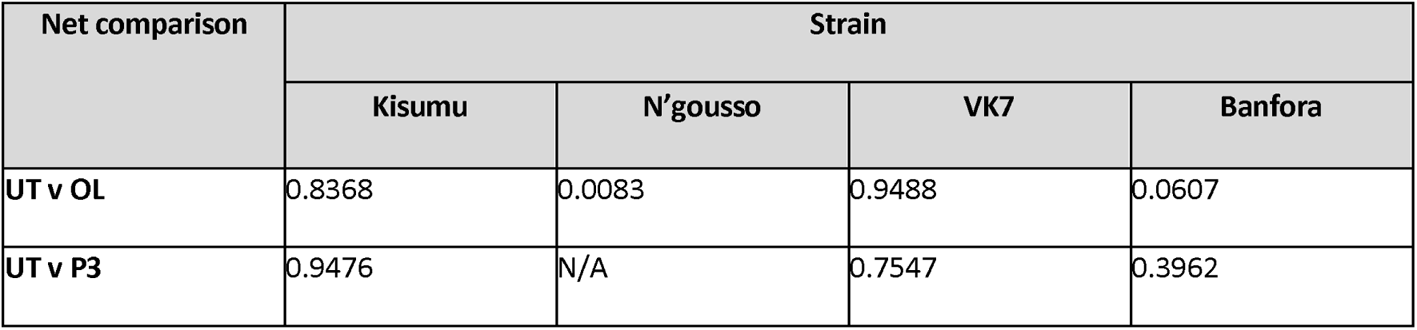
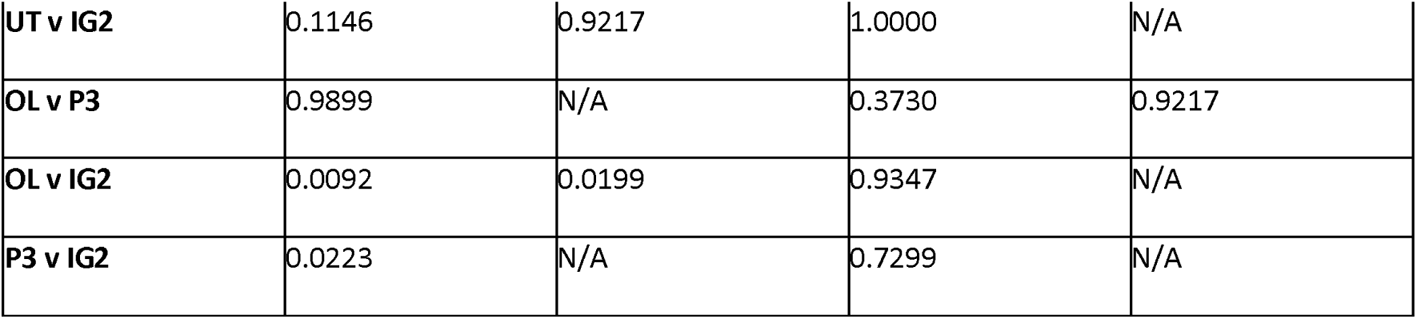
Average contact duration in first 10minutes – within strain, between net comparisons.

**Additional Table 15.**
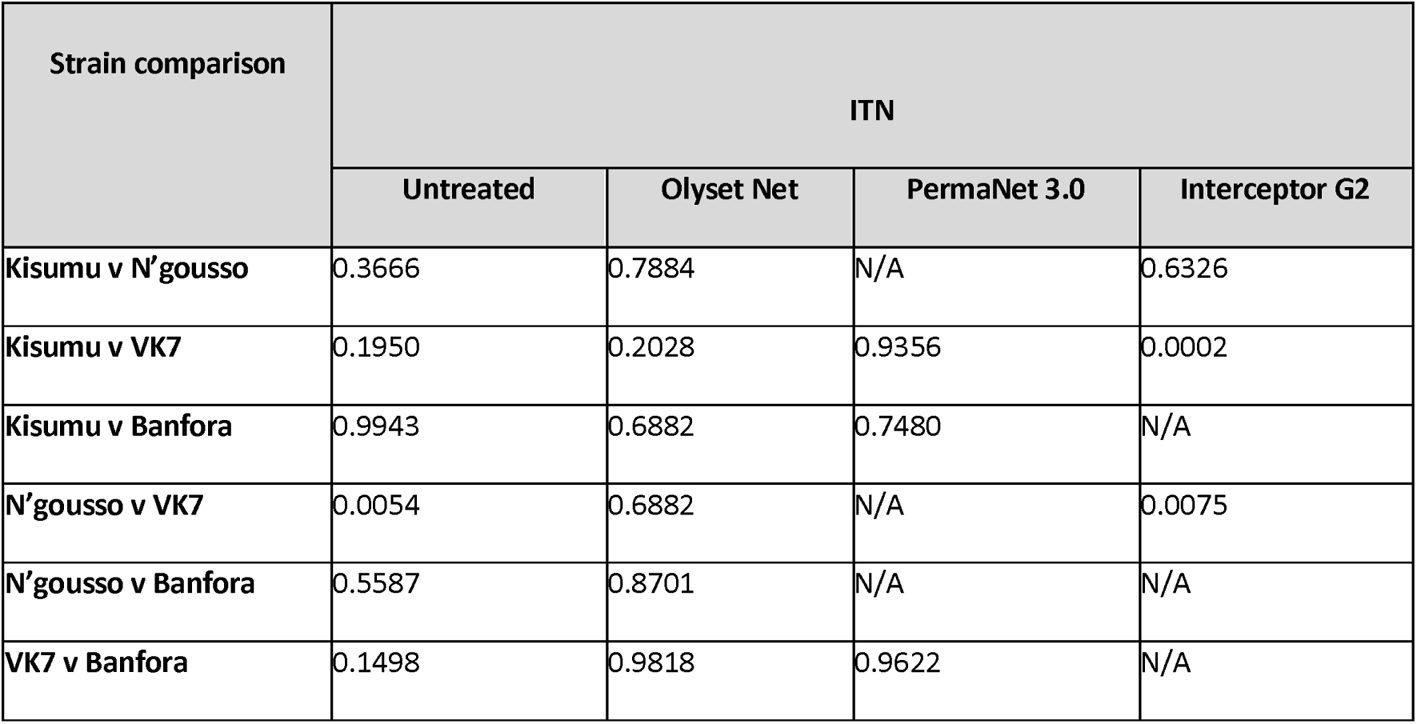
Average contact duration in first 10minutess – within net, between strain comparisons.

**Additional Table 16.**
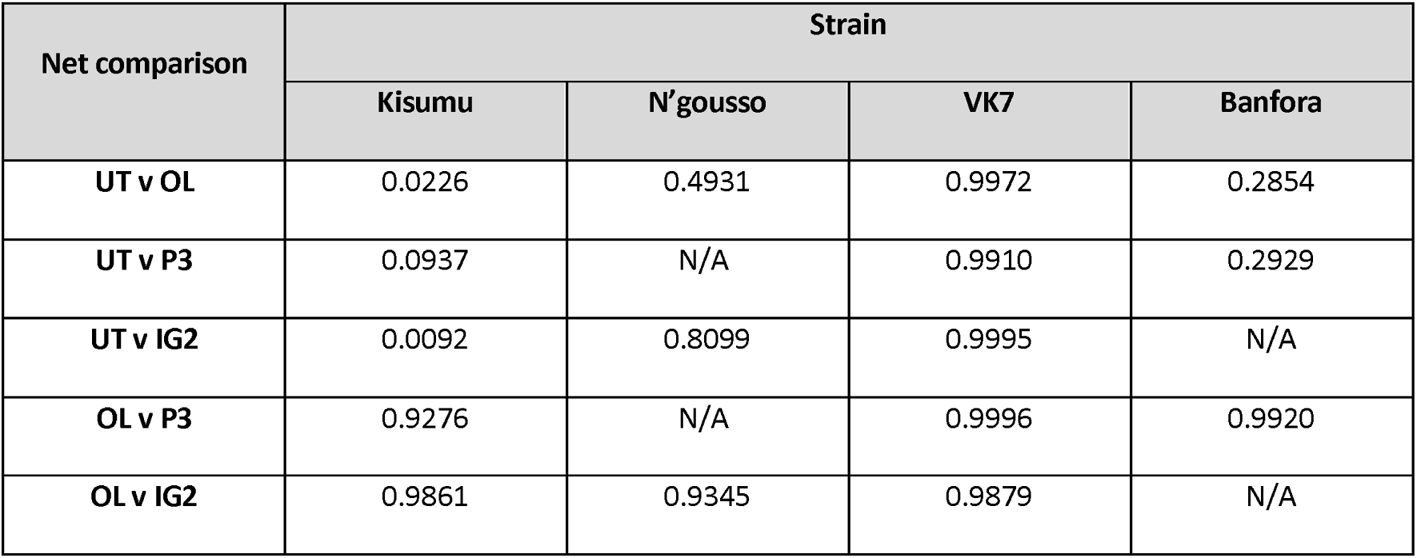
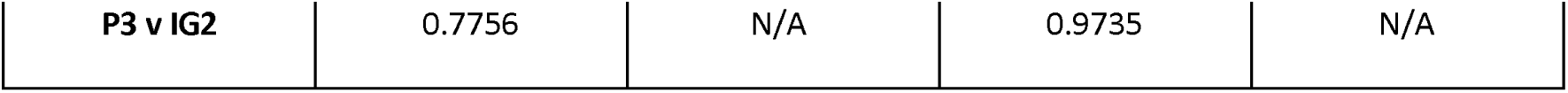
Comparison (p-value) of average swooping speeds across 2hour assay within four different strains, between four different net treatments.

**Additional Table 17.**
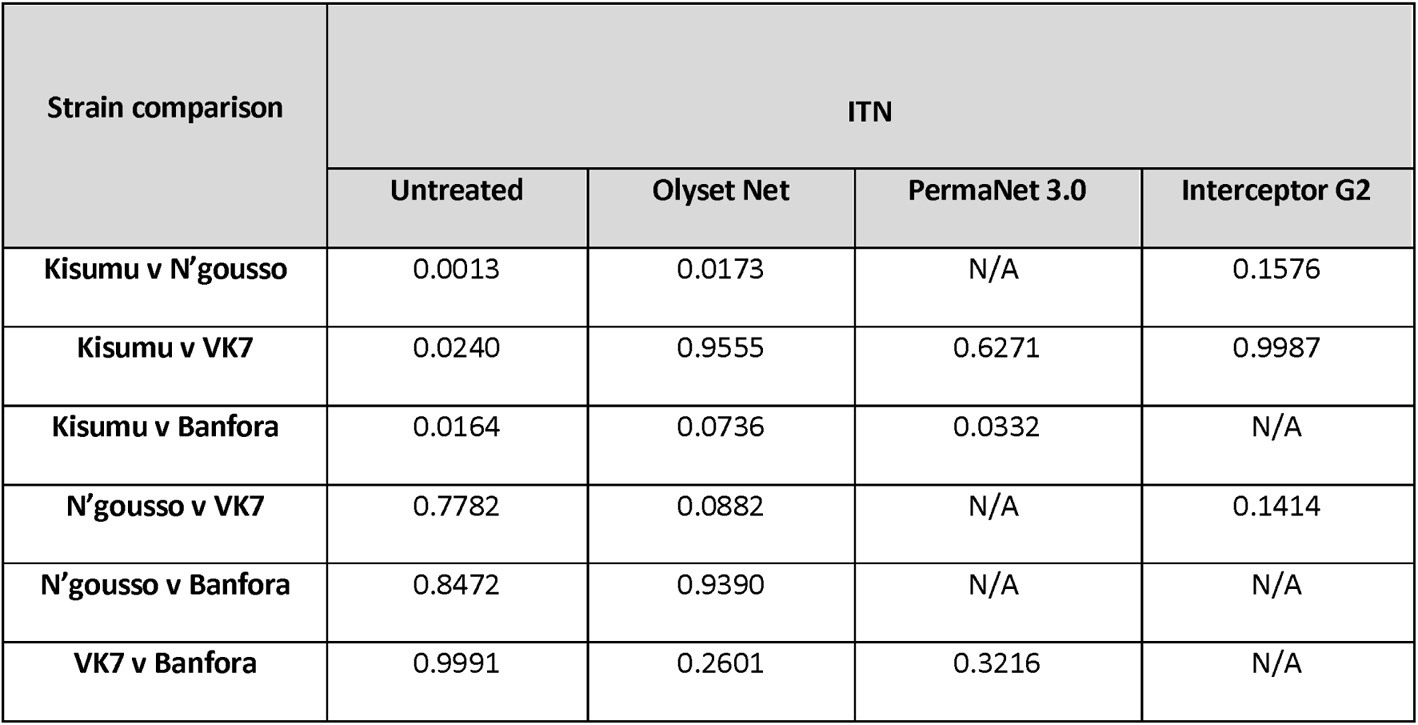
Comparison of average swooping speeds across 2hour assay within four net treatments, between four strains.

**Additional Table 18.**
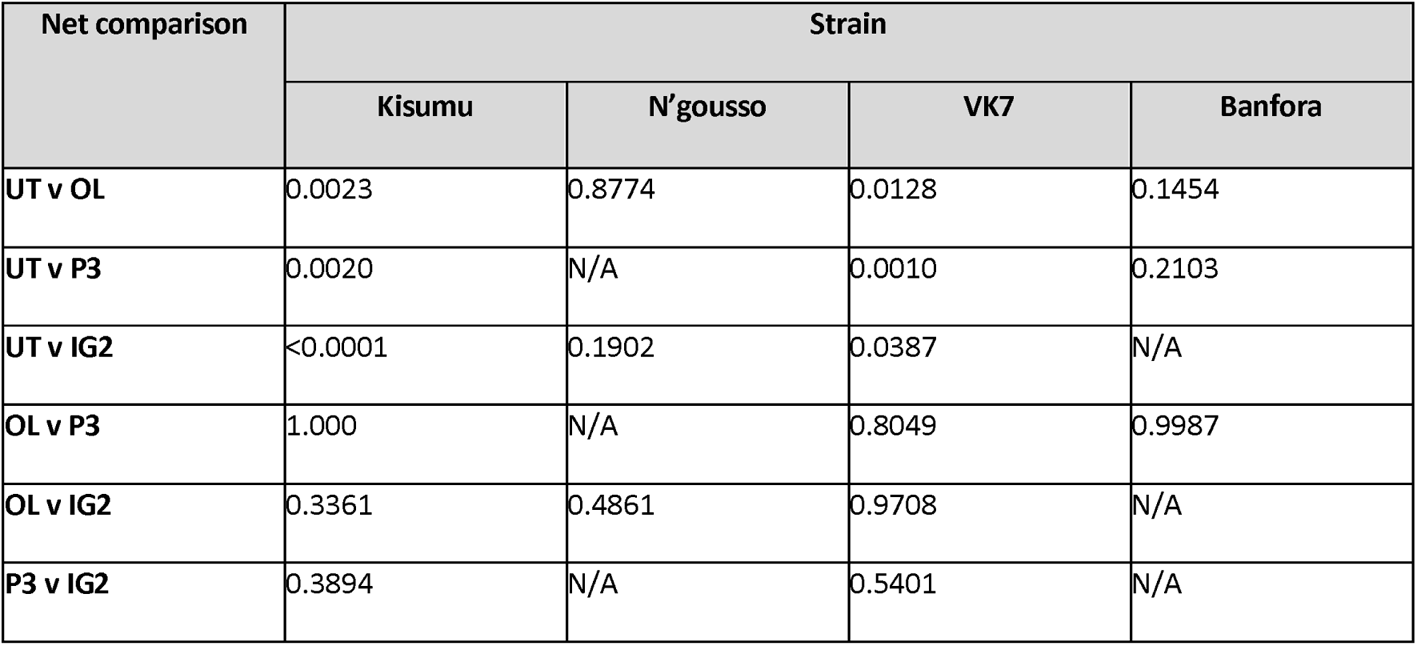
Comparison of activity decay over time (p-value), within strain, between net treatment.

**Additional Table 19.**
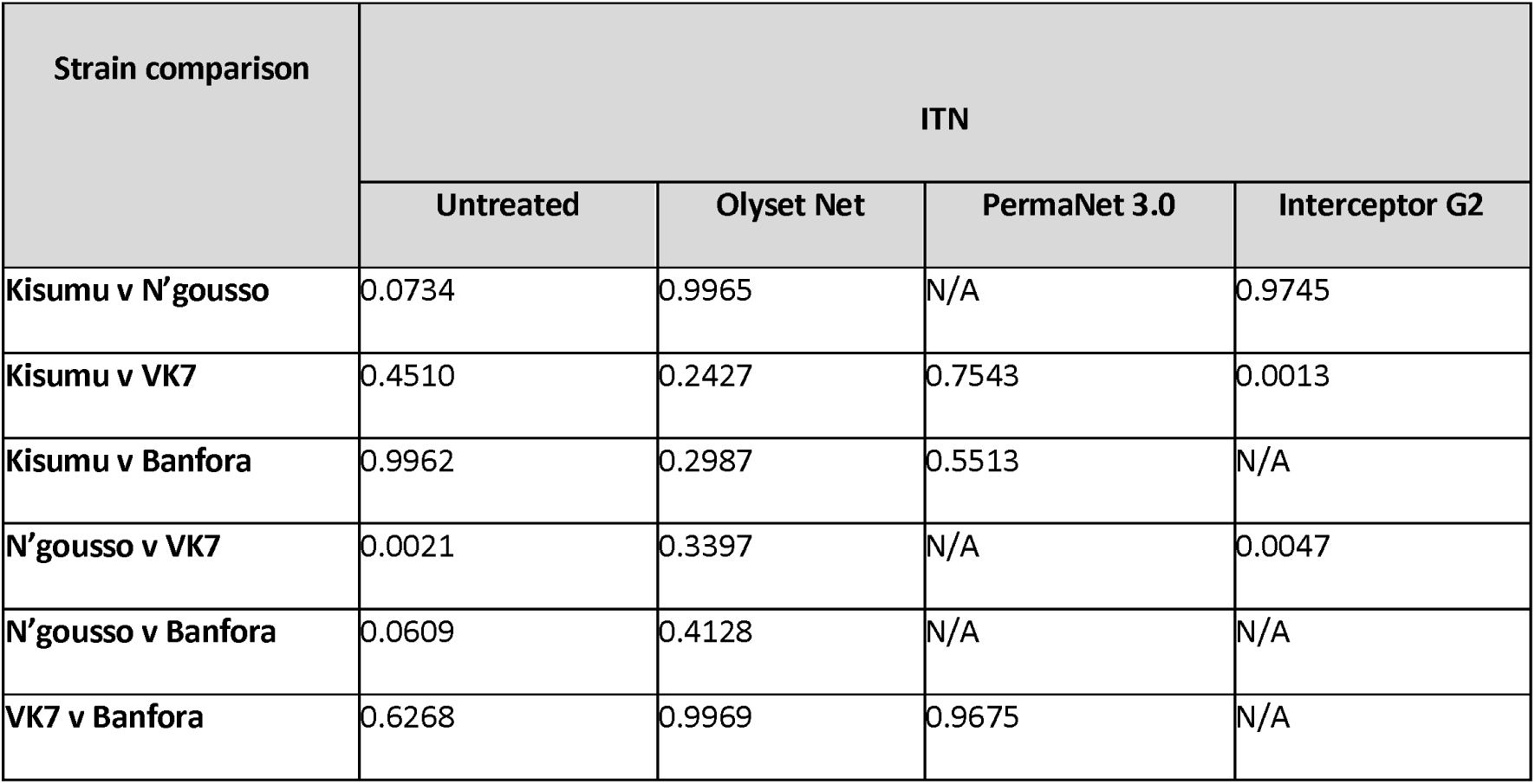
Comparison of activity decay over time (p-value), within net treatment, between strains.

## Notes

### Competing Interest Statement

The authors have declared no competing interest.

