## Supplementary tables for "Behaviour of pyrethroid resistant Anopheles gambiae at the interface of two dual active-ingredient bed nets, assessed by room-scale infrared video tracking"

Additional material

**Additional Table 1. Mean 24hour mortality [95% CI]**

| **Treatment** | **Strain** | **24hour mortality (%) [95% CI]** |
| --- | --- | --- |
| Untreated | Kisumu | 9.5 [1.47, 17,54] |
|  | N’gousso | 17.64 [7.87, 27.40] |
|  | VK7 | 3.36 [0, 10.05] |
|  | Banfora | 4.52 [3.85, 5.20] |
| Olyset Net | Kisumu | 98.67 [95.24, 100] |
|  | N’gousso | 97.97 [94.44, 100] |
|  | VK7 | 20.35 [2.09, 38.01] |
|  | Banfora | 45.34 [14.52, 76.17] |
| PermaNet 3.0 | Kisumu | 100 [100, 100] |
|  | N’gousso | 100 [100, 100] |
|  | VK7 | 71.37 [51.39,91.36] |
|  | Banfora | 72.38 [41.13, 100] |
| Interceptor G2 | Kisumu | 93.88 [81.53, 100] |
|  | N’gousso | 94.56 [91.10, 98.02] |
|  | VK7 | 15.90 [8.62, 23.21] |

**Additional Table 2. Mean 24hour mortality comparisons between three insecticide treated nets and four mosquito strains, two susceptible (Kisumu and N’gousso) and two resistant (VK7 and Banfora).**

| **Strain comparison** | **ITN** | | |
| --- | --- | --- | --- |
|  | **Olyset Net** | **PermaNet 3.0** | **Interceptor G2** |
| Kisumu v VK7 | t(9)= 12.80, p<0.0001 | t(10)= 3.68, p=0.0042 | t(8)= 16.64, p<0.0001 |
| Kisumu v Banfora | t(10)= 4.42, p0.0013 | t(7)= 5.81, p=0.0007 | N/A |
| N’gousso v VK7 | t(9)= 12.67, p<0.0001 | t(6)= 2.07, p=0.0903 | t(8)= 21.54, p<0.0001 |
| N’gousso v Banfora | t(10)= 4.36, p=0.0014 | t(3)= 2.95, p=0.0602 | N/A |

**Additional Table 3. Comparison of median survival times of susceptible (Kisumu and N’gousso) and resistant (VK7 and Banfora) strains on four different net treatments.**

| **Strain comparison** | **ITN** | | | |
| --- | --- | --- | --- | --- |
|  | **Untreated net** | **Olyset Net** | **PermaNet 3.0** | **Interceptor G2** |
| **Kisumu v N’gousso** | χ² (1, N=221) = 6.68, p=0.0098 | χ² (1, N=285) = 0.12, p=0.7241 | χ² (1, N=188) = 0.00, p>0.9999 | χ² (1, N=190) = 0.42, p=0.5191 |
| **Kisumu v VK7** | χ² (1, N=192) = 0.01, p=0.9733 | χ² (1, N=267) = 134.40, p<0.0001 | χ² (1, N=284) = 47.72, p<0.0001 | χ² (1, N=234) = 102.80, p<0.0001 |
| **Kisumu v Banfora** | Χ² (1, N=191) = 2.55, p=0.1102 | χ² (1, N=268) = 67.23, p<0.0001 | χ² (1, N=212) = 43.99, p<0.0001 | N/A |
| **N’gousso v VK7** | χ² (1, N=205) = 7.11, p=0.0077 | χ² (1, N=272) = 133.50, p<0.0001 | χ² (1, N=184) = 15.98, p<0.0001 | χ² (1, N=228) = 124.10, p<0.0001 |
| **N’gousso v Banfora** | χ² (1, N=204) = 15.67, p<0.0001 | χ² (1, N=263) = 65.16, p<0.0001 | χ² (1, N=112) = 14.67, p=0.0001 | N/A |
| **VK7 v Banfora** | χ² (1, N=175) = 3.12, p=0.0773 | χ² (1, N=255) = 3.70, p=0.0545 | χ² (1, N=208) = 3.63, p=0.0568 | N/A |

**Additional Table 4. Statistically significant differences (p values) in total activity time split into four different behavioural modes (swooping, visiting, bouncing and resting), comparing untreated (UT) net to either Olyset Net (OL), PermaNet 3.0 (P3) or Interceptor G2 (IG2), for susceptible (Kisumu and N’gousso) and resistant (VK7 and Banfora) mosquitoes.**

| **Strain** | **Behaviour** | **Insecticide treated net** | | |
| --- | --- | --- | --- | --- |
|  |  | **Olyset Net** | **PermaNet 3.0** | **Interceptor G2** |
| Kisumu | swooping | <0.0001 | 0.0006 | 0.0067 |
|  | visiting | <0.0001 | <0.0001 | <0.0001 |
|  | bouncing | <0.0001 | <0.0001 | <0.0001 |
|  | resting | <0.0001 | <0.0001 | <0.0001 |
| N’gousso | swooping | <0.0001 | N/A | <0.0001 |
|  | visiting | <0.0001 | N/A | <0.0001 |
|  | bouncing | <0.0001 | N/A | <0.0001 |
|  | resting | <0.0001 | N/A | <0.0001 |
| VK7 | swooping | <0.0001 | <0.0001 | <0.0001 |
|  | visiting | <0.0001 | <0.0001 | <0.0001 |
|  | bouncing | <0.0001 | <0.0001 | <0.0001 |
|  | resting | 0.0300 | 0.0264 | 0.1591 |
| Banfora | swooping | <0.0001 | <0.0001 | N/A |
|  | visiting | <0.0001 | <0.0001 | N/A |
|  | bouncing | <0.0001 | <0.0001 | N/A |
|  | resting | <0.0001 | <0.0001 | N/A |

**Additional Table 5. Within strain comparisons (p-value) of total activity time split into four different behavioural modes (swooping, visiting, bouncing and resting) between three ITNs (Olyset Net = OL, PermaNet 3.0 = P3, Interceptor G2 = IG2).**

| **Behaviour** | **ITN comparison** | **Strain** | | | |
| --- | --- | --- | --- | --- | --- |
|  |  | **Kisumu** | **N’gousso** | **VK7** | **Banfora** |
| Swooping | OL v P3 | 0.5855 | N/A | 0.8362 | 0.9811 |
|  | OL v IG2 | 0.1778 | 0.9800 | 0.5898 | N/A |
|  | P3 v IG2 | 0.8577 | N/A | 0.9740 | N/A |
| Visiting | OL v P3 | 0.3119 | N/A | 0.9528 | 0.9015 |
|  | OL v IG2 | 0.2388 | 0.3841 | 0.6678 | N/A |
|  | P3 v IG2 | 0.9985 | N/A | 0.9275 | N/A |
| Bouncing | OL v P3 | 0.1818 | N/A | 0.9674 | 0.8342 |
|  | OL v IG2 | 0.0961 | 0.3402 | 0.6559 | N/A |
|  | P3 v IG2 | 0.9897 | N/A | 0.8978 | N/A |
| Resting | OL v P3 | 0.2797 | N/A | 0.9999 | 0.2268 |
|  | OL v IG2 | 0.3265 | 0.5880 | 0.8416 | N/A |
|  | P3 v IG2 | 0.9997 | N/A | 0.8657 | N/A |

**Additional Table 6. Within treatment comparisons (p-value) of total activity split into for behavioural modes (swooping, visiting, bouncing and resting) on four ITNs (Untreated net = UT, Olyset Net = OL, PermaNet 3.0 = P3, Interceptor G2 = IG2) between four mosquito strains**

| **Behaviour** | **Strain comparison** | **ITN** | | | |
| --- | --- | --- | --- | --- | --- |
|  |  | **UT** | **OL** | **P3** | **IG2** |
| Swooping | Kisumu v N’gousso | 0.0950 | 0.4483 | N/A | 0.9978 |
|  | Kisumu v VK7 | 0.0010 | 0.4572 | 0.6651 | 0.8395 |
|  | Kisumu v Banfora | 0.0640 | 0.0879 | 0.9477 | N/A |
|  | N’gousso v VK7 | 0.4166 | 0.9999 | N/A | 0.7475 |
|  | N’gousso v Banfora | 0.9984 | 0.8000 | N/A | N/A |
|  | VK7 v Banfora | 0.5157 | 0.8484 | 0.9748 | N/A |
| Visiting | Kisumu v N’gousso | 0.0352 | 0.7484 | N/A | 0.9844 |
|  | Kisumu v VK7 | 0.0248 | 0.6266 | 0.9997 | 0.9377 |
|  | Kisumu v Banfora | 0.5026 | 0.7043 | 0.9989 | N/A |
|  | N’gousso v VK7 | 0.9994 | 0.9946 | N/A | 0.9997 |
|  | N’gousso v Banfora | 0.5523 | 0.9998 | N/A | N/A |
|  | VK7 v Banfora | 0.4796 | 0.9980 | 0.9999 | N/A |
| Bouncing | Kisumu v N’gousso | 0.3291 | 0.9915 | N/A | 0.7701 |
|  | Kisumu v VK7 | <0.0001 | 0.9997 | 0.5171 | 0.7801 |
|  | Kisumu v Banfora | 0.0014 | 0.9985 | 0.8952 | N/A |
|  | N’gousso v VK7 | 0.0032 | 0.9822 | N/A | 1.000 |
|  | N’gousso v Banfora | <0.0001 | 0.9697 | N/A | N/A |
|  | VK7 v Banfora | <0.0001 | 0.9999 | 0.9669 | N/A |
| Resting | Kisumu v N’gousso | 0.1368 | 0.5673 | N/A | 0.3099 |
|  | Kisumu v VK7 | 0.0004 | 1.0000 | 0.3601 | 0.8771 |
|  | Kisumu v Banfora | 0.8988 | 0.9949 | 0.8801 | N/A |
|  | N’gousso v VK7 | 0.0668 | 0.5917 | N/A | 0.7962 |
|  | N’gousso v Ban | 0.1368 | 0.3588 | N/A | N/A |
|  | VK7 v Banfora | 0.0001 | 0.9891 | 0.1704 | N/A |

**Additional Table 7. Mean total number of bed net contacts [95% CI], mean total contact duration [95% CI] and maximum number of mosquitoes seen in one frame of video recording.**

| **ITN** | **Strain** | **Replicates** | **Mean total number of contacts [95% CI]** | **Mean total contact duration [95% CI]** | **Maximum number of mosquitoes** |
| --- | --- | --- | --- | --- | --- |
| UT | Kisumu | 5 | 74885  [53016.58, 96753.42]] | 9044.20  [7723.32, 10202.13] | 29 |
|  | N’gousso | 4 | 62162.25  [39731.46, 84593.04] | 8254.66  [5049.89, 11459.43] | 17 |
|  | VK7 | 4 | 41811.25  [31737, 51885.5] | 5783.19  [3589.78, 7976.59] | 14 |
|  | Banfora | 4 | 80824  [43804.85, 117843.20] | 11005.31  [7866.99, 14143.62] | 14 |
| OL | Kisumu | 6 | 6169.17  [2521.05, 9817.29] | 622.03  [238.49, 1005.57] | 16 |
|  | N’gousso | 5 | 4531.17  [3175.86, 5886.47] | 342.37  [256.65, 428.09] | 10 |
|  | VK7 | 6 | 7393.2  [3465.90, 11320.50] | 682.53  [532.04, 833.02] | 9 |
|  | Banfora | 6 | 7413.5  [3695.42, 11167.58] | 787.19  [268.22, 1306.16] | 14 |
| P3 | Kisumu | 6 | 10909.33  [3149.47, 18669.20] | 1929.23  [187.19, 3671.27] | 27 |
|  | VK7 | 5 | 6219  [2576.16, 9861.84] | 1164.04  [[736.89, 1591.20] | 9 |
|  | Banfora | 3 | 14772  [1453.14, 28090.86] | 1668.92  586.24, 2751.60] | 8 |
| IG2 | Kisumu | 6 | 12759.5  [6312.24, 19206.76] | 2236.16  [1434.93, 3037.40] | 14 |
|  | N’gousso | 6 | 6686.5  [2384.28, 10988.75] | 1822.44  [971.85, 2673.03] | 16 |
|  | VK7 | 5 | 10488  [4366.25, 16609.75] | 1587.40  [853.88, 2320.90] | 11 |

**Additional Table 8. Within strain statistical comparisons (p value) of total number of net contacts for susceptible (Kisumu and N’gousso) and resistant (VK7 and Banfora) mosquitoes between ~~three ITNs~~four nets (UT = untreated, OL = Olyset Net, P3 = PermaNet 3.0, IG2 = Interceptor G2).**

| **Net comparison** | **Strain** | | | |
| --- | --- | --- | --- | --- |
|  | **Kisumu** | **N’gousso** | **VK7** | **Banfora** |
| **UT v OL** | <0.0001 | <0.0001 | <0.0001 | <0.0001 |
| **UT v P3** | <0.0001 | N/A | <0.0001 | <0.0001 |
| **UT v IG2** | <0.0001 | <0.0001 | <0.0001 | N/A |
| **OL v P3** | 0.7873 | N/A | 0.9966 | 0.6402 |
| **OL v IG2** | 0.5684 | 0.9741 | 0.9445 | N/A |
| **P3 v IG2** | 0.9833 | N/A | 0.8689 | N/A |

**Additional Table 9. Within treatment statistical comparisons (p value) of total number of net contacts for four nets between four mosquito strains.**

| **Strain comparison** | **ITN** | | | |
| --- | --- | --- | --- | --- |
|  | **Untreated** | **Olyset Net** | **PermaNet 3.0** | **Interceptor G2** |
| **Kisumu v N’gousso** |  | 0.9883 | N/A | 0.6322 |
| **Kisumu v VK7** | <0.0001 | 0.9957 | 0.8151 | 0.9738 |
| **Kisumu v Banfora** |  | 0.9948 | 0.9250 | N/A |
| **N’gousso v VK7** | 0.0095 | 0.9496 | N/A | 0.8914 |
| **N’gousso v Banfora** | 0.0202 | 0.9414 | N/A | N/A |
| **VK7 v Banfora** | <0.0001 | 1.0000 | 0.5473 | N/A |

**Additional Table 10. Within strain comparisons (p-value) of total duration of net contact for susceptible (Kisumu and N’gousso) and resistant (VK7 and Banfora) mosquitoes between three ITNs (OL = Olyset Net, P3 = PermaNet 3.0, IG2 = Interceptor G2).**

| **Net comparison** | **Strain** | | | |
| --- | --- | --- | --- | --- |
|  | **Kisumu** | **N’gousso** | **VK7** | **Banfora** |
| **UT v OL** | <0.0001 | <0.0001 | <0.0001 | <0.0001 |
| **UT v P3** | <0.0001 | N/A | <0.0001 | <0.0001 |
| **UT v IG2** | <0.0001 | <0.0001 | <0.0001 | <0.0001 |
| **OL v P3** | 0.1265 | N/A | 0.8889 | 0.6018 |
| **OL v IG2** | 0.0373 | 0.0617 | 0.5123 | N/A |
| **P3 v IG2** | 0.9514 | N/A | 0.9088 | N/A |

**Additional Table 11. Within treatment comparison (p-value) of total net contact duration for three ITNs between four mosquito strains.**

| **Strain comparison** | **ITN** | | | |
| --- | --- | --- | --- | --- |
|  | **Untreated** | **Olyset Net** | **PermaNet 3.0** | **Interceptor G2** |
| **Kisumu v N’gousso** |  | 0.9567 | N/A | 0.8908 |
| **Kisumu v VK7** | 0.0001 | 0.9994 | 0.5914 | 0.7097 |
| **Kisumu v Banfora** | 0.0252 | 0.9938 | 0.9829 | N/A |
| **N’gousso v VK7** | 0.0051 | 0.9310 | N/A | 0.9801 |
| **N’gousso v Banfora** | 0.0015 | 0.8683 | N/A | N/A |
| **VK7 v Banfora** | <0.0001 | 0.9992 | 0.9006 | N/A |

**Additional Table 12. Percentage of contact duration in first the 10minutes of room scale tracking assay – within strain, between net differences.**

| **Net comparison** | **Strain** | | | |
| --- | --- | --- | --- | --- |
|  | **Kisumu** | **N’gousso** | **VK7** | **Banfora** |
| **UT v OL** | <0.0001 | <0.0001 | 0.9999 | 0.9965 |
| **UT v P3** | 0.0121 | N/A | 0.0547 | 0.8800 |
| **UT v IG2** | 0.0003 | 0.0108 | 0.9592 | N/A |
| **OL v P3** | 0.0626 | N/A | 0.0312 | 0.9302 |
| **OL v IG2** | 0.5533 | 0.0243 | 0.9327 | N/A |
| **P3 v IG2** | 0.6108 | N/A | 0.1253 | N/A |

**Additional Table 13. Percentage of contact duration in first 10mins of assay – within net, between strain differences**

| **Strain comparison** | **ITN** | | | |
| --- | --- | --- | --- | --- |
|  | **Untreated** | **Olyset Net** | **PermaNet 3.0** | **Interceptor G2** |
| **Kisumu v N’gousso** | 0.9829 | 0.7099 | N/A | 0.9489 |
| **Kisumu v VK7** | 0.9717 | <0.0001 | 0.8614 | 0.0004 |
| **Kisumu v Banfora** | 0.9996 | <0.0001 | 0.1913 | N/A |
| **N’gousso v VK7** | 0.8703 | <0.0001 | N/A | 0.0021 |
| **N’gousso v Banfora** | 0.9707 | <0.0001 | N/A | N/A |
| **VK7 v Banfora** | 0.9884 | 0.9062 | 0.5609 | N/A |

**Additional Table 14. Average contact duration in first 10minutes – within strain, between net comparisons.**

| **Net comparison** | **Strain** | | | |
| --- | --- | --- | --- | --- |
|  | **Kisumu** | **N’gousso** | **VK7** | **Banfora** |
| **UT v OL** | 0.8368 | 0.0083 | 0.9488 | 0.0607 |
| **UT v P3** | 0.9476 | N/A | 0.7547 | 0.3962 |
| **UT v IG2** | 0.1146 | 0.9217 | 1.0000 | N/A |
| **OL v P3** | 0.9899 | N/A | 0.3730 | 0.9217 |
| **OL v IG2** | 0.0092 | 0.0199 | 0.9347 | N/A |
| **P3 v IG2** | 0.0223 | N/A | 0.7299 | N/A |

**Additional Table 15. Average contact duration in first 10minutess – within net, between strain comparisons.**

| **Strain comparison** | **ITN** | | | |
| --- | --- | --- | --- | --- |
|  | **Untreated** | **Olyset Net** | **PermaNet 3.0** | **Interceptor G2** |
| **Kisumu v N’gousso** | 0.3666 | 0.7884 | N/A | 0.6326 |
| **Kisumu v VK7** | 0.1950 | 0.2028 | 0.9356 | 0.0002 |
| **Kisumu v Banfora** | 0.9943 | 0.6882 | 0.7480 | N/A |
| **N’gousso v VK7** | 0.0054 | 0.6882 | N/A | 0.0075 |
| **N’gousso v Banfora** | 0.5587 | 0.8701 | N/A | N/A |
| **VK7 v Banfora** | 0.1498 | 0.9818 | 0.9622 | N/A |

**Additional Table 16. Comparison (p-value) of average swooping speeds across 2hour assay within four different strains, between four different net treatments.**

| **Net comparison** | **Strain** | | | |
| --- | --- | --- | --- | --- |
|  | **Kisumu** | **N’gousso** | **VK7** | **Banfora** |
| **UT v OL** | 0.0226 | 0.4931 | 0.9972 | 0.2854 |
| **UT v P3** | 0.0937 | N/A | 0.9910 | 0.2929 |
| **UT v IG2** | 0.0092 | 0.8099 | 0.9995 | N/A |
| **OL v P3** | 0.9276 | N/A | 0.9996 | 0.9920 |
| **OL v IG2** | 0.9861 | 0.9345 | 0.9879 | N/A |
| **P3 v IG2** | 0.7756 | N/A | 0.9735 | N/A |

**Additional Table 17. Comparison of average swooping speeds across 2hour assay within four net treatments, between four strains.**

| **Strain comparison** | **ITN** | | | |
| --- | --- | --- | --- | --- |
|  | **Untreated** | **Olyset Net** | **PermaNet 3.0** | **Interceptor G2** |
| **Kisumu v N’gousso** | 0.0013 | 0.0173 | N/A | 0.1576 |
| **Kisumu v VK7** | 0.0240 | 0.9555 | 0.6271 | 0.9987 |
| **Kisumu v Banfora** | 0.0164 | 0.0736 | 0.0332 | N/A |
| **N’gousso v VK7** | 0.7782 | 0.0882 | N/A | 0.1414 |
| **N’gousso v Banfora** | 0.8472 | 0.9390 | N/A | N/A |
| **VK7 v Banfora** | 0.9991 | 0.2601 | 0.3216 | N/A |

**Additional Table 18. Comparison of activity decay over time (p-value), within strain, between net treatment.**

| **Net comparison** | **Strain** | | | |
| --- | --- | --- | --- | --- |
|  | **Kisumu** | **N’gousso** | **VK7** | **Banfora** |
| **UT v OL** | 0.0023 | 0.8774 | 0.0128 | 0.1454 |
| **UT v P3** | 0.0020 | N/A | 0.0010 | 0.2103 |
| **UT v IG2** | <0.0001 | 0.1902 | 0.0387 | N/A |
| **OL v P3** | 1.000 | N/A | 0.8049 | 0.9987 |
| **OL v IG2** | 0.3361 | 0.4861 | 0.9708 | N/A |
| **P3 v IG2** | 0.3894 | N/A | 0.5401 | N/A |

**Additional Table 19. Comparison of activity decay over time (p-value), within net treatment, between strains.**

| **Strain comparison** | **ITN** | | | |
| --- | --- | --- | --- | --- |
|  | **Untreated** | **Olyset Net** | **PermaNet 3.0** | **Interceptor G2** |
| **Kisumu v N’gousso** | 0.0734 | 0.9965 | N/A | 0.9745 |
| **Kisumu v VK7** | 0.4510 | 0.2427 | 0.7543 | 0.0013 |
| **Kisumu v Banfora** | 0.9962 | 0.2987 | 0.5513 | N/A |
| **N’gousso v VK7** | 0.0021 | 0.3397 | N/A | 0.0047 |
| **N’gousso v Banfora** | 0.0609 | 0.4128 | N/A | N/A |
| **VK7 v Banfora** | 0.6268 | 0.9969 | 0.9675 | N/A |
